# Influenza H3 and H1 hemagglutinins have different genetic barriers for resistance to broadly neutralizing stem antibodies

**DOI:** 10.1101/2019.12.30.891135

**Authors:** Nicholas C. Wu, Andrew J. Thompson, Juhye M. Lee, Wen Su, Britni M. Arlian, Jia Xie, Richard A. Lerner, Hui-Ling Yen, Jesse D. Bloom, Ian A. Wilson

**Author notes:** These authors contributed equally to this work (Nicholas C. Wu, Andrew J. Thompson). To whom correspondence may be addressed. (I.A.W.).

## Abstract

In the past decade, the discovery and characterization of broadly neutralizing antibodies (bnAbs) to the highly conserved stem region of influenza hemagglutinin (HA) have provided valuable insights for development of a universal influenza vaccine. However, the genetic barrier for resistance to stem bnAbs has not been thoroughly evaluated. Here, we performed a series of deep mutational scanning experiments to probe for resistance mutations. We found that the genetic barrier to resistance to stem bnAbs is generally very low for the H3 subtype but substantially higher for the H1 subtype. Several resistance mutations in H3 cannot be neutralized by stem bnAbs at the highest concentration tested, do not reduce *in vitro* viral fitness and *in vivo* pathogenicity, and are often present in circulating strains as minor variants. Thus, H3 HAs have a higher propensity than H1 HAs to escape major stem bnAbs and creates a potential challenge in the development of a *bona fide* universal influenza vaccine.

**ONE SENTENCE SUMMARY:** Acquisition of resistance by influenza virus to broadly neutralizing hemagglutinin stem antibodies varies tremendously depending on subtype.

## Introduction

The major surface antigen of influenza virus, the hemagglutinin (HA), is composed of a highly variable globular head domain that houses the receptor binding site and a conserved stem domain that is responsible for membrane fusion (*1*). All of the major antigenic sites on HA are located on the HA globular head (*2*–*5*), which is immunodominant over the stem (*6*). However, most antibodies to the globular head domain are strain-specific. In contrast, although harder to elicit during natural infection or vaccination, many HA stem antibodies have impressive cross-reactive breadth (*7*, *8*). The isolation, characterization and structure determination of broadly neutralizing antibodies (bnAbs) to the HA stem over the past decade have provided tremendous insights into antiviral and vaccine development against influenza virus (*9*), including immunogen design towards a universal influenza vaccine (*10*–*12*). Several stem bnAbs are also currently in clinical trials as therapeutics (*13*). Stem bnAbs have also provided templates for design of small proteins, peptides and small molecules against influenza virus (*14*–*18*). Therefore, while influenza virus remains a major global health concern, stem bnAbs open up multiple promising avenues to tackle this challenging problem.

However, emergence of resistance mutations can be a major obstacle for antiviral and vaccine development. Several studies have reported difficulty in selecting strong resistance mutations to stem bnAbs even after extensive passaging of the viruses (*19*–*21*), or through deep mutational scanning (*22*), which is a comprehensive and unbiased approach (*23*). Nonetheless, strong resistance mutations have been reported in other studies through virus passaging (*20*, *24*, *25*). It is unclear then why some studies were able to identify strong resistance mutations while others could not. Here we systematically compare how readily resistance can emerge to stem bnAbs in H3 and H1 HAs, and find that there are major differences between the subtypes.

### Deep mutational scanning of the major HA stem epitope

CR9114 (*26*) and FI6v3 (*27*) are two bnAbs that bind the HA stem and have exceptional neutralization breadth. They are in fact two of the known bnAbs with the greatest breadth against influenza viruses. Both FI6v3 and CR9114 neutralize group 1 and 2 influenza A viruses (*26*, *27*), and CR9114 further cross-reacts with influenza B HA (*26*). Deep mutational scanning (*23*), which combines saturation mutagenesis and next-generation sequencing, has previously been applied to study how HA mutations affect influenza viral fitness (*28*–*30*), and to identify viral mutants that are resistant to anti-HA antibodies (*31*). Here, we employed deep mutational scanning of the HA stem on influenza virus to search for resistance mutations to CR9114 and FI6v3. We focused on eight HA2 residues in the HA stem of H3N2 A/Hong Kong/1/1968 (H3/HK68): namely Q42, I45, D46, Q47, I48, N49, L52, and T111 (Fig. 1, A to C). All except T111 are located on HA2 helix A, which is a common target for stem bnAbs. Residues 42, 45, 46, 48, 49, 52 interact with CR9114 and FI6v3, whereas residue 47 only interacts with FI6v3 (Fig. 1, D and E). The completely buried T111 was also selected because its mutation in H5 HA enabled escape from CR6261 (*24*), which binds a similar epitope to CR9114 (*26*, *32*).

**Fig. 1.**
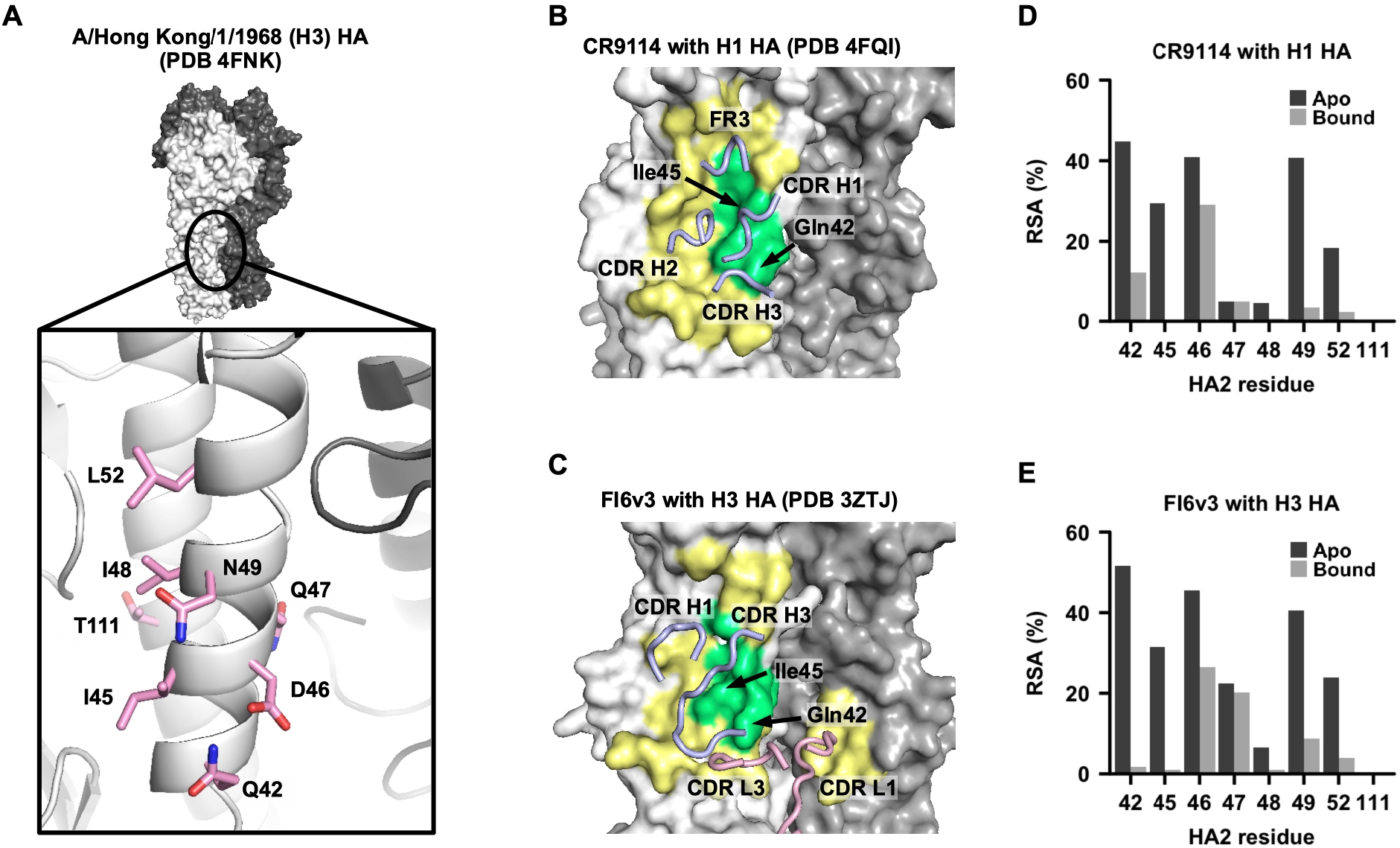
Epitopes of broadly neutralizing antibodies to the HA stem. (**A**) The location of residues of interest in this study on the HA structures. All residues of interest are on HA2. One protomer of the trimer is shown in light gray and the other two protomers in dark gray and a detailed view of the location of the residues of interest is shown in the inset. (**B-C**) Epitopes of (**B**) CR9114 Fab in complex with H1 HA (PDB 4FQI) (*26*) and (**C**) FI6v3 in complex with H3 HA (PDB 3ZTJ) (*27*) are colored in yellow and green, and residues of interest colored in green. The arrows indicate the positions of HA2 residues 42 and 45, which are in the center of the bnAb epitopes. Antibody paratopes (CDRs and FR regions) are shown in tube representation and labeled accordingly. Blue: heavy chain. Pink: light chain. (**D-E**) The relative solvent accessibility (RSA) of each residue of interest is shown. Black bar: apo form. Gray bar: Fab-bound form.

We quantified the *in vitro* fitness of 147 out of 152 possible single viral mutants and 6,234 out of 10,108 possible double viral mutants across the eight residues of interest in H3/HK68 HA2 under five different conditions: no antibody, 2 μg/mL CR9114 IgG, 10 μg/mL CR9114 IgG, 0.3 μg/mL FI6v3 IgG, and 2.5 μg/mL FI6v3 IgG (fig. S1). In the absence of antibody, many viral mutants have a relative fitness [proxy for replication fitness (*30*)], similar to wild type (WT), which was set as 1 (Fig. 2, A and B), and indicate that the HA stem region can tolerate many mutations.

**Fig. 2.**
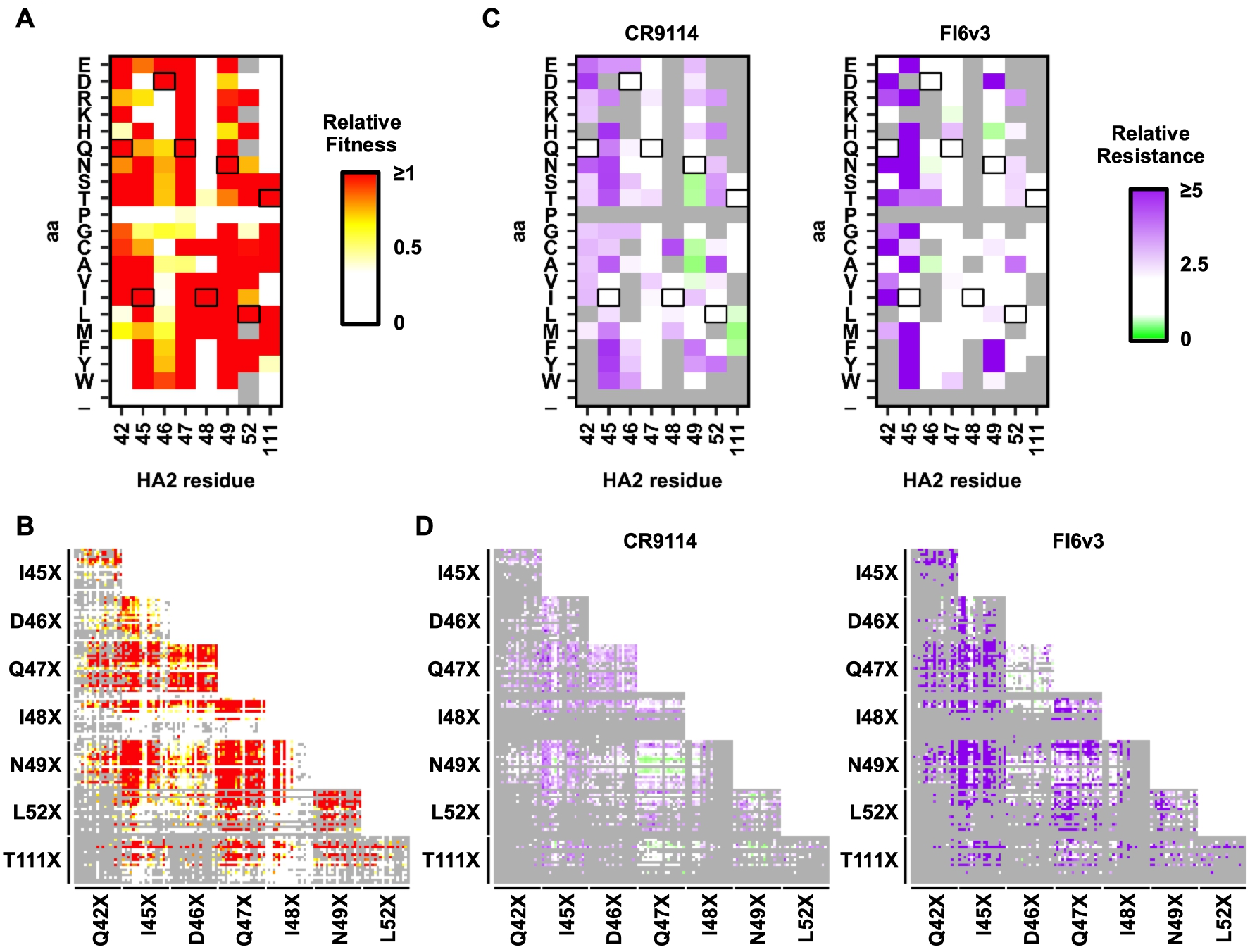
Fitness and resistance profile of H3/HK68 HA2 single and double viral mutants. (**A-B**) Based on the deep mutational scanning experiment, the relative fitness of (**A**) single and (**B**) double mutants are shown with wild type (WT) set as 1. In (**A**) and (**B**), mutants with a next-generation sequencing read count of less than 20 from the plasmid mutant library are excluded and shown as grey. (**C-D**) Relative resistance for (**C**) each single and (**D**) each double mutant against 10 μg/mL CR9114 antibody or 2.5 μg/mL FI6v3 antibody is shown. Relative resistance for WT is set as 1. In (**C**) and (**D**), mutants with a relative fitness of less than 0.5 are shown as grey. Residues correspond to WT sequence are boxed. In (**B**) and (**D**), each color point in the heatmap represents a double mutant. Of note, some mutants with an increased sensitivity to antibody are shown in green on the relative resistance color scale.

We further quantified the relative resistance of each viral mutant by normalizing their relative fitness in the presence and absence of antibody (fig. S2). Many resistance mutations to CR9114 and FI6v3 were observed (Fig. 2, C and D) and most were located at HA2 residues 42 and 45, which form an important component of the binding interface with CR9114 and FI6v3 (Fig. 1, D and E). In addition, the double mutants also showed high relative resistance if one mutation exhibited high relative resistance even if the other did not (fig. S3). Overall, these results demonstrate the prevalence of H3/HK68 resistance mutations to stem bnAbs and mutations with cross-resistance to both CR9114 and FI6v3, even though these bnAbs are encoded by different germline genes and have very different angles of approach to the HA (*8*).

### Validation of resistance mutations

To validate our findings from deep mutational scanning, 24 single and double HA mutants of the H3/HK68 virus that spanned a range of relative resistance were individually constructed and tested against CR9114 and FI6v3 IgGs (Fig. 3A). The minimum inhibitory concentration (MIC) in a microneutralization assay strongly correlated with the relative resistance from deep mutational scanning (Spearman’s rank correlation > 0.8, Fig. 3B) with several viral mutants showing strong cross-resistance to both CR9114 and FI6v3. For example, the MICs of CR9114 and FI6v3 against mutants I45Y/S/N/F/W were all >100 and ≥20 μg mL^−1^, respectively, compared to 3.1 and 0.2 μg mL^−1^ for WT. This validation experiment substantiates our finding that strong resistance mutations are prevalent in H3/HK68.

**Fig. 3.**
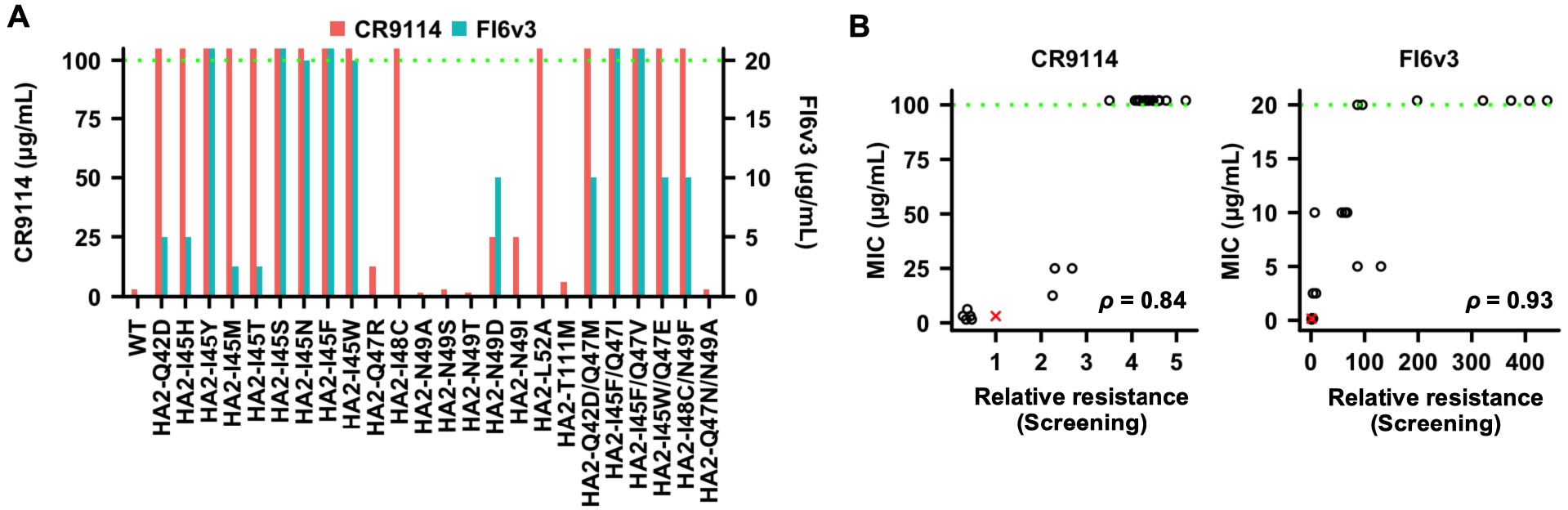
Characterization of antibody-resistant mutants. (**A**) The minimum inhibitory concentration (MIC) of CR9114 and FI6v3 to individual viral mutants are shown. The MIC of CR9114 is in red and represented by the y-axis on the left. The MIC of FI6v3 is in blue and represented by the y-axis on the right. (**B**) The Spearman’s rank correlations (*ρ*) between the MIC measured from individual mutants and the relative resistance (against 10 μg/mL CR9114 antibody or 2.5 μg/mL FI6v3 antibody) computed from the profiling experiment (screening) are shown. The green dashed line represents the upper detection limit in **A** and **B.** Wild type is represented by the red “X”.

### Natural occurrence of resistance mutations

Next, we explored whether these resistance mutations were found in naturally circulating strains While most strong resistance mutations have not yet been observed in naturally circulating strains, it is important to note that a few could be identified at low frequency in natural human H3N2 isolates (*33*), including I45T, I45M and N49D (Fig. 4A). I45T is also observed in human H3N2 isolates sequenced without any passaging (fig. S4A), implying that its presence was not due to a passaging artifact (*34*). Moreover, the strong cross-resistance mutation I45F was found in all human H2N2 viruses that circulated from 1957 to 1968 (Fig. 4B, fig. S4B), while almost all avian H2N2 viruses have Ile45 (Fig. 4C), and explains why it is more difficult for human H2N2 viruses to be bound or neutralized by some stem bnAbs compared to other subtypes (*24*, *26*, *35*). Thus, these findings suggest that some resistance mutations to stem bnAbs already occur in circulating strains.

**Fig. 4.**
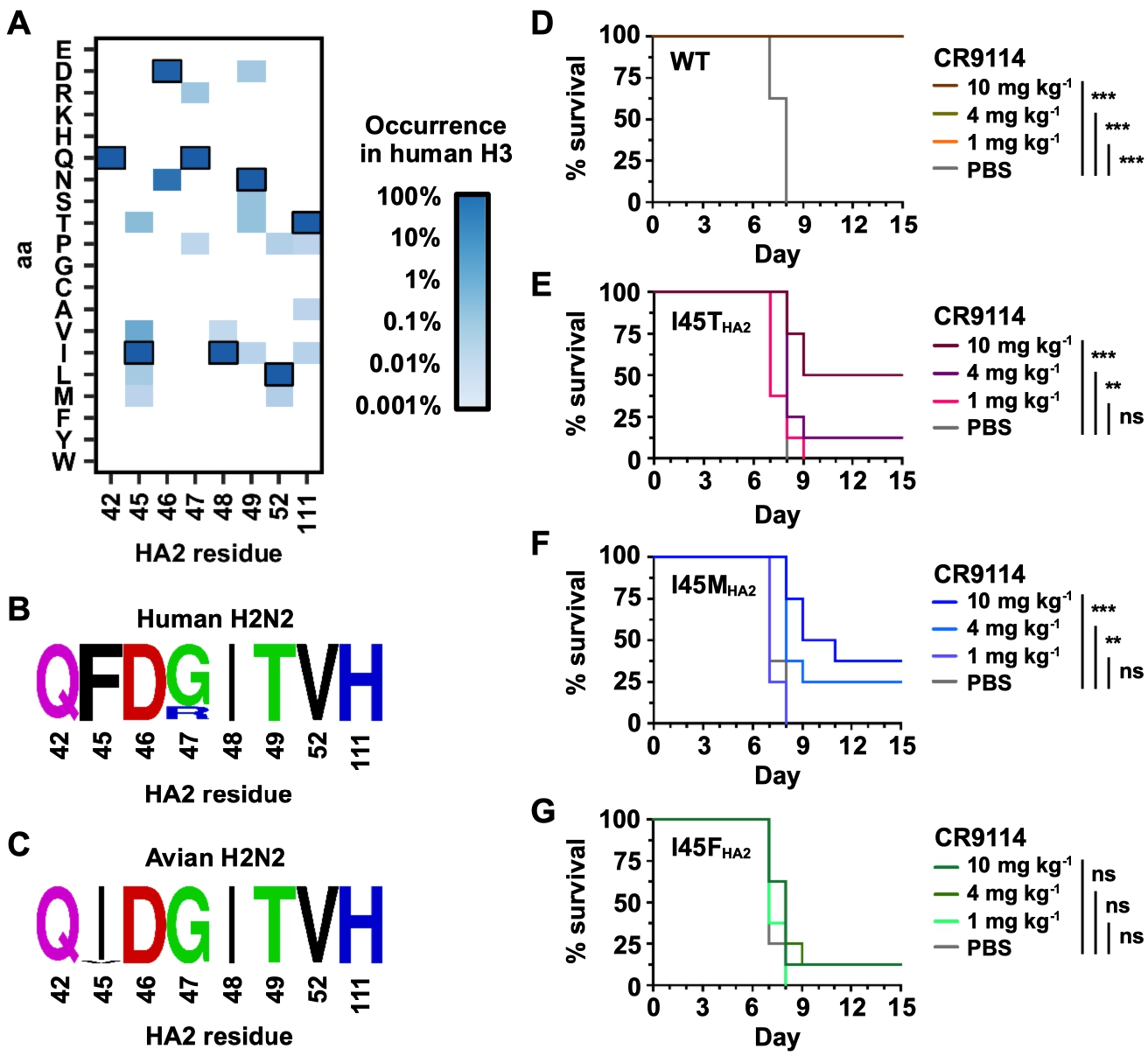
*In vivo* characterization and natural occurrence of antibody-resistant mutants. (**A**) The natural occurrence frequencies of different amino-acid variants at the residues of interest in human H3 HAs are shown as a heatmap. (**B-C**) The natural occurrence frequencies of different amino-acid variants at the residues of interest in (**B**) human H2N2 HA or (**C**) avian H2N2 HA are shown as sequence logos. (**D-G**) Prophylactic protection experiments were performed with different doses of CR9114 against (**D**) WT, (**E**) HA2 I45T mutant, (**F**) HA2 I45M mutant, and (**G**) HA2 I45F mutant. Recombinant H3/HK68 (7:1 on H1/PR8 backbone) viruses were used. Lethal doses (25 mLD_50_) of WT or mutant viruses were used. Kaplan-Meier survival curves are shown. Paired analysis of each treatment group, relative to control, was conducted using Log-rank (Mantel-Cox) tests. *** indicates p-value ≤ 0.001; ** indicates p-value ≤ 0.01; * indicates p-value ≤ 0.05; ns (not significant) indicates p-value > 0.05.

### *In vivo* pathogenicity and escape of resistance mutations

We further tested the *in vivo* viral pathogenicity of HA2 mutants I45T, I45M, and I45F, which are of relevance to circulating strains (see above). The weight loss profiles in mice after infection by HA2 mutant and WT viruses were comparable (fig. S5), indicating that these resistance mutations do not reduce *in vivo* pathogenicity. We further demonstrated that I45T, I45M, and I45F were able to escape *in vivo* prophylactic protection. While mice infected with WT were completely protected by CR9114 IgG at all tested doses (1, 4, and 10 mg kg^−1^), mutants I45T, I45M, and I45F were lethal even at the highest dose of CR9114 IgG (Fig. 4, D to G).

### Resistance mutations decrease affinity to bnAbs

To dissect the resistance mechanism, we tested the binding of H3/HK68 I45T, I45M, and I45F recombinant HAs to CR9114 and FI6v3, and also to another stem bnAb 27F3 (*35*), which utilizes the same V_H_1-69 germline as CR9114 and similarily neutralizes group 1 and 2 influenza A viruses. The binding (K_d_) of CR9114 Fab, CR9114 IgG, 27F3 Fab, 27F3 IgG, and FI6v3 IgG was all diminished against the HA mutants compared to WT (Table 1 and fig. S6), and was particularly dramatic with the I45F mutant, where binding was undetectable to CR9114 Fab and IgG, and 27F3 Fab and IgG. In contrast, the binding of these stem Fabs and IgGs to N49T, which did not exhibit any resistance against CR9114 and FI6v3 (Fig. 2C, Fig. 3A), are comparable to the WT (Table 1). As a control, we also tested binding of bnAb S139/1 that targets the receptor-binding site far from the stem epitope (*36*, *37*). S139/1 IgG affinities against those HA mutants (K_d_ = 1.8 nM to 3.1 nM) were similar to WT (K_d_ = 2.1 nM). Thus, virus resistance to stem bnAbs correlated with a decrease in binding affinity to the mutant HAs.

**Table 1.** Binding affinity of IgGs and Fabs against HAs of WT and different H3/HK68 HA2 mutants.

| **K<sub>d</sub> in nM** | **WT** | **HA2-I45T** | **HA2-I45M** | **HA2-I45F** | **HA2-N49T** |
| --- | --- | --- | --- | --- | --- |
| **CR9114 Fab** | 43.2 ± 0.4 | 779.6 ± 50.0 | 319.1 ± 12.1 | n.b. | 11.3 ± 0.4 |
| **27F3 Fab** | 122.6 ± 3.0 | n.b. | n.b. | n.b. | 105.0 ± 1.0 |
| **CR9114 IgG** | < 0.1 | 164.8 ± 2.3 | 75.0 ± 1.0 | n.b. | < 0.1 |
| **27F3 IgG** | 1.7 ± 0.1 | n.b. | n.b. | n.b. | 1.9 ± 0.9 |
| **FI6v3 IgG** | < 0.1 | 7.8 ± 0.2 | 6.7 ± 0.1 | 62.0 ± 0.2 | < 0.1 |
| **S139/1 IgG** | 2.1 ± 0.1 | 3.1 ± 0.3 | 1.9 ± 0.1 | 1.8 ± 0.1 | 2.8 ± 0.1 |

471 n.b.: no binding

To understand the structural basis of the resistance, we determined crystal structures of HAs with HA2 mutations I45T, I45M, and I45F to 2.1 to 2.5 Å resolutions (table S1 and fig. S7A). Compared to WT (Ile45), the shorter side chain of I45T would create a void when CR9114 is bound (fig. S7B) that would be energetically unfavorable. In contrast, the longer flexible side chain of I45M would likely clash with CR9114 (fig. S7B), but CR9114 is still able to bind the I45M mutant, albeit with much lower affinity than WT (Table 1). The I45F mutant, however, makes a more severe clash with CR9114 and no binding was detected (Table 1, fig. S7B). Similar observations for FI6v3 (fig. S7C) explain the sensitivity of CR9114 and FI6v3 to mutations at HA2 residue 45.

### Resistance to HA stem bnAbs is subtype specific

We further aimed to examine whether those mutations that confered strong resistance in H3/HK68 would have the same phenotypes in other H3 strains. Consequently, we examined the phenotype of three resistance mutations of relevance to circulating strains (see below), namely I45T, I45M, and I45F, on an H3N2 A/Wuhan/359/95 (H3/Wuhan95) genetic background. These three viral mutants had WT-like titer after viral rescue and passaging (fig. S8A), showed similar plaque size as WT (fig. S8B), and conferred strong resistance to FI6v3 (fig. S8, C and D). Therefore, these mutants have similar phenotypes in H3/HK68 and H3/Wuhan95 viruses. This result led us to hypothesize that strong resistance mutants to stem bnAbs are readily attainable in a wide range of H3 strains, as well as to explore whether the same phenomenon can be observed in H1 subtype, which is the other currently circulating influenza A subtype in the human population.

Previously, two of the authors here performed deep mutational scanning to search for resistance mutants of H1N1 A/WSN/33 (H1/WSN) virus to FI6v3 (*22*). In contrast to this study, resistance mutations to FI6v3 were rare in H1/WSN, and had only very small effects. This discrepancy suggested that the prevalence of resistance mutations is markedly different between H3/HK68 and H1/WSN. We therefore performed four additional deep mutational scanning experiments – three with H1N1 strains, namely A/Solomon Islands/3/2006 (H1/SI06) against FI6v3, A/Michigan/45/2015 (H1/Mich15) against FI6v3, H1/WSN against CR9114, and one with H3N2 strain A/Perth/16/2009 (H3/Perth09) against FI6v3. The H1/SI06 and H1/Mich15 HA mutant virus libraries contain all possible single amino-acid substitutions at HA2 residues 42, 45, 46, 47, 48, 49, 52, and 111, whereas H1/WSN and H3/Perth09 HA mutant virus libraries both contain all possible single substitutions across the entire HA and were constructed in previous studies (*38*, *39*). We also analyzed the previously published dataset on H1/WSN against FI6v3 (*22*).

To compare H1/SI06, H1/Mich15, H1/WSN and H3/Perth09 to H3/HK68, we computed the relative resistance of mutations at HA2 residues 42, 45, 46, 47, 48, 49, 52, and 111 (Fig. 5, A to E). Similar to H3/HK68, resistance mutations are highly prevalent in H3/Perth09 (Fig. 5C). Conversely, resistance mutations were rare in H1/SI06 (Fig. 5A), H1/Mich15 (Fig. 5B), and H1/WSN (Fig. 5, D and E). We further calculated the *fraction surviving* (*22*) for each viral mutant across the entire H1/WSN and H3/Perth09 HA proteins during antibody selection. Fraction surviving is a quantitative measure for the resistance that is normalized across deep mutational scanning experiments (*22*). The fraction surviving values of H1/WSN mutants against CR9114 were all very small, similar to previous observations of H1/WSN against FI6v3 (Fig. 5F, fig. S10). In stark contrast, many mutants of H3/Perth09 were identified with a large fraction surviving value (Fig. 5F). Consistent with the relative resistance profile of H3/HK68 (Fig. 2C), a number of H3/Perth09 mutants with a large fraction surviving value were again at HA2 residues 42 and 45 (Fig. 5F and fig. S11A). Moreover, mutations at HA2 residue 53, which were not examined in H3/HK68 (Fig. 2), had high fraction surviving in H3/Perth09 against FI6v3 (fig. S11, A and B). In H3 HA, mutation of HA2 residue 53 would abolish a hydrogen bond to the complementarity-determining region (CDR) H3 of FI6v3 (fig. S11B). Together, these results suggest that the prevalence of resistance mutations to stem bnAbs is a general phenomenon for the H3 subtype, but not the H1 subtype.

**Fig. 5.**
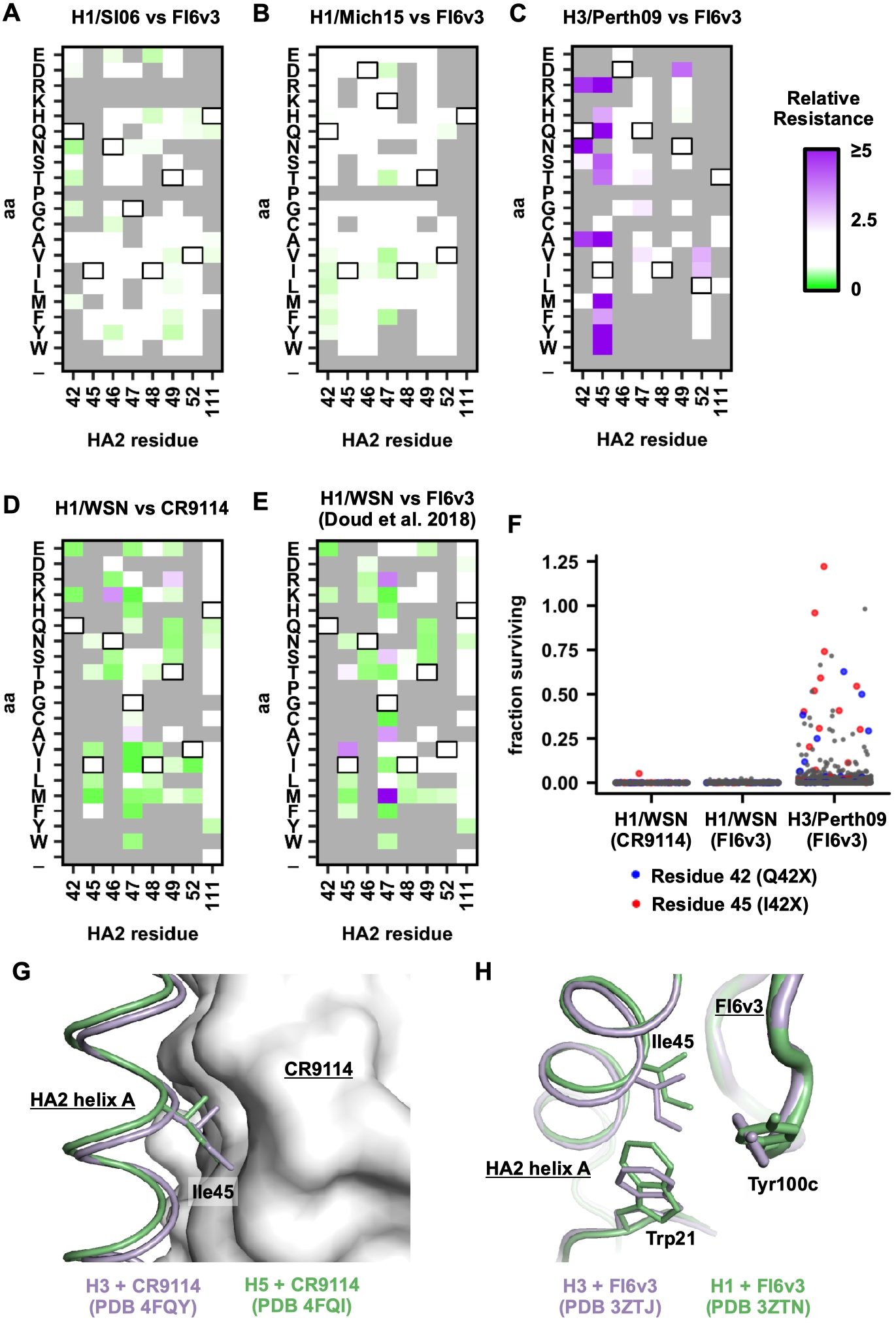
Relative resistance profile of single mutants in multiple strains of influenza virus. (**A-E**) Relative resistance is measured for each single viral mutant at HA2 residues 42, 45, 46, 27, 48, 49, 52, and 111 of (**A**) H1/SI06 against 300 ng/mL FI6v3 antibody, (**B**) H1/Mich15 against 300 ng/mL FI6v3 antibody, (**C**) H1/WSN against 100 ng/mL CR9114 antibody, (**D**) H1/WSN against 200 ng/mL FI6v3 antibody (data are from (*22*)), and (**E**) H3/Perth09 against 15 μg/mL FI6v3 antibody. Relative resistance for WT is set as 1. Mutants with a relative fitness of less than 0.5 are excluded and shown as grey. Residues correspond to WT sequence are boxed. (**F**) Fraction surviving for all single mutants across the HA protein are shown. Each data point represents one mutant. Fraction surviving was computed as previously described (*22*). Assuming no antibody-mediated enhancement of virus replication, the theoretical upper limit for the fraction surviving is 1, which indicates that the replication fitness is the same with and without antibody selection (in practice, fraction surviving values slightly > 1 can sometimes be obtained due to the experimental error in qPCR and next-generation sequencing). Data for H1/WSN against FI6v3 were from a previous study (*22*). Data points that represent mutations at residues 42 and 45 are colored in blue and red, respectively, and mutations at other residues are in grey. (**G**) The crystal structures of H3 HA in complex with CR9114 (PDB 4FQY) (*26*) and H5 HA in complex with CR9114 (PDB 4FQI) (*26*) were compared by aligning their CR9114 heavy chain variable domains. (**H**) Similarly, H3 HA in complex with FI6v3 (PDB 3ZTJ) (*27*) and H1 HA in complex with FI6v3 (PDB 3ZTN) (*27*) were compared by aligning their heavy chain variable domains.

### Subtype-specific differences in the HA stem

We next aimed to elucidate the mechanism that underlies the lower genetic barrier to resistance to stem bnAbs in H3 HA as compared to H1 HA. Many mutations at HA2 residue 45 have a high fitness cost in H1/SI06 (fig. S9A), which can increase the genetic barrier to resistance. However, most mutations at HA2 residue 45 have no fitness cost in H1/Mich15. In addition, the mutational fitness profiles of H1/SI06 and H1/Mich15 (fig. S9, A and B) show that many mutations can be tolerated in the HA stem, similar to H3/HK68 (Fig. 2A, fig. S9, C to F). Thus, the difference in genetic barrier to resistance to stem bnAbs between H1 and H3 subtypes cannot be fully explained by their ability to tolerate mutations (i.e. fitness cost of mutations).

We therefore further compared the structures of CR9114 in complex with H3 HA and in complex with H5 HA (Fig. 5G) (*26*). Since the structure of CR9114 with H1 HA is not available, CR9114 with H5 HA was used instead, as it also belongs to group 1 HAs and is therefore more similar to H1 than to H3 HA (group 2). Structural comparison indicates that CR9114 packs tighter to the helix A of H3 HA than to H5 HA. Specifically, there is ~1 Å difference in the position of the Cα of HA2 Ile45. Subsequently, a bulkier substitution at HA2 Ile45, such as I45M, would create a larger disruption of the CR9114-HA binding interface in the context of H3 subtype. Thus, subtle differences in the binding of bnAbs to different HA subtypes may lead to differences in how antibodies are affected by mutations in or near the epitope.

Similar observations can be made for FI6v3. The orientation of Tyr100c on CDR H3 of FI6v3 differs when binding to H1 or H3 HAs (Fig. 5H) (*27*). The position of HA2 Ile45 also differs between H1 and H3 HAs when FI6v3 is bound. As a result, Tyr100c of FI6v3 packs tighter to HA2 Ile45 of H3 than to H1 HA. Thus, a bulkier substitution at HA2 Ile45 will disrupt binding between FI6v3 and H3 HA to a greater extent than FI6v3 to H1 HA. Therefore, the low genetic barrier to resistance to stem bnAbs in the H3 subtype can be at least partly attributed to both high mutational tolerance in the HA stem and subtype-specific structural features. While a number of subtype-specific structural features are known in the stem region (*40*), how these structural differences influence the genetic barriers for resistance to stem bnAbs remains to be addressed in future studies.

### Ramifications for escape from a universal vaccine or therapeutic stem bnAbs

Prior studies of influenza bnAbs have not considered whether different subtypes might have different abilities to generate resistance mutations against bnAbs. A major finding here is that H3 HA has a much lower genetic barrier to resistance to two of the broadest bnAbs, CR9114 and FI6v3, as compared to H1 HA. This observation is consistent with the literature, where strong resistance to other human HA stem antibodies have been reported in H3 subtype (*20*, *25*, *41*) versus none (*20*) to weak resistance (*21*, *22*) in the H1 subtype. Therefore, it may be easier for stem bnAbs to maintain suppression of the H1 subtype than the H3 subtype.

Since the HA stem is immunosubdominant to the globular head domain, immunological pressure on the HA stem may not have been sufficient to impact the evolution of circulating influenza strains (*42*). However, several stem bnAbs are currently in clinical trials for therapeutic purposes (*13*) and a number of recently developed influenza vaccine immunogens have focused on targeting the HA stem (*8*, *9*). If stem bnAbs begin to be distributed on a global scale, the immunological pressure on the HA stem will certainly surge to a level not previously seen. Our findings here indicate that resistance mutations could emerge, at least in H3 subtype.

Although resistance mutations to stem bnAbs are still rare in currently circulating influenza strains (Fig. 4A), it is important to evaluate the potential impact of such mutations since many vaccine strategies aim to elicit anti-stem antibodies. In fact, we were not able to overcome some key resistance mutations (I45T, I45M, and I45F) by *in vitro* evolution of CR9114 (fig. S12). Nonetheless, the best strategy to prevent or overcome such resistance may involve delivery or elicitation of a combination of antibodies with different resistance profiles. In addition, it remains to be explored whether stem bnAbs exist or can be generated that are difficult to escape from the H3 subtype. The discovery and characterization of bnAbs with different escape profiles will therefore continue to be key to broaden our arsenal against influenza virus. For example, human H2N2 virus, which carries a Phe at HA2 residue 45, often has low reactivity with stem bnAbs (*24*, *26*, *27*, *35*, *43*), although a very few can have high potency against human H2N2 (*44*–*46*). Future studies on anti-stem responses against human H2N2 and emerging viruses, such as H5N1 and H7N9, may provide further insights into how to overcome potential resistance when immune pressure is transferred to the HA stem.

## ACKNOWLEDGEMENTS

We thank Wenli Yu and Geramie Grande for technical support in protein expression, Steven Head, Jessica Ledesma and Padmaja Natarajan at TSRI Next Generation Sequencing Core and Fred Hutch Genomics Core for next-generation sequencing, Matthew Haynes and Brian Seegers of the TSRI Flow Cytometry Core Facility for performing FACS, Alfred Ho, Eva-Maria Strauch, and Barney Graham for insightful discussions, and James Paulson for his continual support.

## Author Contributions

N.C.W., J.M.L., R.A.L., H.L.Y., J.D.B., and I.A.W. conceived and designed the study. N.C.W. and J.M.L. performed the deep mutational scanning experiments. N.C.W., J.M.L., and J.D.B. performed the computational data analysis. N.C.W. performed the structural analysis and yeast display experiment. N.C.W. and W.S. performed functional characterization of the viral mutants. A.J.T. and B.M.A. performed the *in vivo* characterization of the viral mutants. N.C.W. and J.X. produced the CR9114 and 27F3 antibodies. J.M.L. produced the FI6v3 antibody. N.C.W. and I.A.W. wrote the paper and all authors reviewed and edited the paper.

## Data and materials availability

Raw sequencing data will be deposited to the NIH Short Read Archive prior to publication. The X-ray coordinates and structure factors will be deposited to the RCSB Protein Data Bank prior to publication. Links to computer codes and processed data are in supplementary materials. All the other data that support the conclusions of the study are available from the corresponding author upon request.

## SUPPLEMENTARY MATERIALS

Materials and Methods

Figs. S1 to S12

Tables S1 to S3

References 47-66

