## Supplementary Materials for "Influenza H3 and H1 hemagglutinins have different genetic barriers for resistance to broadly neutralizing stem antibodies"

**This PDF file includes:**

Materials and Methods

Figs. S1 to S12

Tables S1 to S3

References 47-66

### Materials and Methods

#### Cell cultures

HEK293T cells and MDCK-SIAT1 cells (Sigma-Aldrich) were maintained in DMEM medium (Thermo Fisher Scientific) supplemented with 10% fetal bovine serum (Thermo Fisher Scientific), 1x MEM non-essential amino acids (Thermo Fisher Scientific) and 100 U $mL^{-1}$ of Penicillin-Streptomycin (Thermo Fisher Scientific). Madin-Darby Canine Kidney (MDCK) cells and Vero cell lines were maintained in minimal essential medium (MEM) containing 10% fetal bovine serum and 1% Penicillin-Streptomycin. Human embryonic kidney 293T cells were maintained in Opti-MEM I reduced serum media containing 5% fetal bovine serum and 1% Penicillin-Streptomycin. MDCK-SIAT1-TMPRSS2 cells (*39*) were maintained in DMEM supplemented with 10% heat-inactivated fetal bovine serum, 2 mM L-glutamine, 100 U $mL^{-1}$ of Penicillin, and 100 $\mu g\ mL^{-1}$ of Streptomycin. Sf9 cells (ATCC) and High Five cells (Thermo Fisher Scientific) were maintained in HyClone insect cell culture medium (GE Healthcare), and Expi293F cells (Thermo Fisher Scientific) in Expi293 expression medium (Thermo Fisher Scientific).

#### Influenza virus

<u>H3N2 A/Hong Kong/1/1968 (H3/HK68)</u>

H3/HK68 virus was generated using a chimeric approach as described previously where the HA and NA were derived from from H3N2 A/Hong Kong/68 virus and the other influenza proteins from H1N1 A/WSN/33 virus (*30*). Briefly, chimeric H3/HK68 HA was constructed based on the pHW2000 plasmid (*47*) that encodes the HA ectodomain ($1_{HA1}$ to $175_{HA2}$, H3 numbering) from A/Hong Kong/1/1968 flanked by the 32-nucleotide 3' non-coding region plus the coding region for the 19 amino acids of the signal peptide from H1/WSN HA at the N terminus, and the coding region for the 46 amino acids of the transmembrane domain and cytoplasmic tail plus 48-nucleotide 5' non-coding region from H1/WSN HA at the C terminus (*30*). For the NA segment, the entire coding region of full-length N1/WSN NA was replaced with that of N2/HK68 NA, with the non-coding regions from N1/WSN. For virus rescue experiments, transfection was performed in HEK293T/MDCK-SIAT1 cells co-culture (ratio of 6:1) using lipofectamine 2000 (Thermo Fisher Scientific) according to the manufacturer's instructions. Virus rescue experiments for H3/HK68 (WT, mutant, or mutant libraries) were performed with the HA and NA segments described above, and the other six WT gene segments from H1/WSN. At 24 hours post-transfection, cells being washed twice with PBS (Thermo Fisher Scientific) and cell culture media were replaced with OPTI-MEM medium (Thermo Fisher Scientific) supplemented with 0.8 μg $mL^{-1}$ TPCK-trypsin (Thermo Fisher Scientific). Virus was harvested at 72 hours post-transfection. MDCK-SIAT1 cells were used for titering and infection, cells were washed twice with PBS prior to the addition of virus, and OPTI-MEM medium supplemented with 0.8 μg $mL^{-1}$ TPCK-trypsin was used.

To generate recombinant H3/HK68 (7:1 on H1/PR8 backbone) viruses, chimeric H1/PR8-flanked H3/HK68 HA was constructed based on the pHW2000 plasmid (*47*) that encodes the HA ectodomain ($1_{HA1}$ to $175_{HA2}$, H3 numbering) from A/Hong Kong/1/1968 flanked by the 32-nucleotide 3' non-coding region and the coding region for the 19 amino acids of the signal peptide from H1/PR8 HA at the N terminus, and the coding region for the 46 amino acids of the transmembrane domain and cytoplasmic tail plus the 48-nucleotide 5' non-coding region from H1/PR8 HA at the C terminus. Virus rescue experiments for H3/HK68 (7:1 on H1/PR8 backbone) viruses (WT or mutant) were the same as described above for chimeric H3/HK68 virus, except that the HA from chimeric H1/PR8-flanked H3/HK68 and the other seven WT segments from H1/PR8 were used.

<u>H3N2 A/Wuhan/359/95 (H3/Wuhan95)</u>

The eight gene segments of the H3/Wuhan95 virus were cloned into the dual promoter pHW2000 vector as described (*48, 49*). Recombinant H3/Wuhan95 viruses were generated in 293T cells using TransIT-LT1 (Mirus). Viruses were passaged twice in MDCK cells containing 1 μg/ml TPCK-trypsin at a multiplicity of infection (MOI) of 0.001 and 0.005, respectively. The HA genes of the recombinant viruses after two passages in MDCK cells were RT-PCR amplified and verified by Sanger sequencing.

**Antibodies**

CR9114 Fab, 27F3 Fab, and 27F3 IgG were expressed as previously described (*26, 35*). For CR9114 IgG expression, the CR9114 heavy and light chains were cloned into pFUSE-CHIg-hG1 and pFUSE2-CLIg-hK respectively. The plasmids were co-transfected into Expi293F cells at 2:1 ratio (light to heavy) using lipofactamine 2000 (Thermo Fisher Scientific) according to manufacturer's instructions. The supernatant was collected at 72 hours post-transfection. All expressed Fabs and IgGs did not contain any affinity tag. Full-length IgG proteins were purified from the supernatant using a protein G column on an ÄKTA$^{TM}$ start (GE Healthcare Life Sciences).

**Construction of H3/HK68 HA mutant libraries**

The HA plasmid mutant libraries were created by ligating a mutant library insert and a PCR-generated vector. The mutant libraries were built based on the pHW2000 plasmid (*47*) that encoded the chimeric A/Hong Kong/1/1968 (H3/HK68) HA (as described above). Silent mutations were introduced at the codons encoding for HA2 residues 43, 44, 50, and 112. These silent mutations acted as an internal barcode to indicate which codon position was being randomized and allowed us to distinguish randomized codon positions from sequencing errors (*50*). The names and nucleotide sequences of individual primers that were used in this study for mutant library construction are listed in table S2.

All PCR reactions were performed using KOD DNA polymerase (EMD Millipore) with 1.5 mM $MgSO_4$, 0.2 mM of each dNTP (dATP, dCTP, dGTP, and dTTP), and 0.6 $\mu$M of forward and reverse primer according to the manufacturer's instructions. All PCR products were purified by gel extraction using PCR Clean-Up and Gel Extraction Kit (Clontech Laboratories).

Insert for the single mutant library

The single mutant library was generated by two PCRs using the WT plasmid as template. In the first PCR, primers StemLib-42-F, StemLib-45-F, StemLib-46-F, StemLib-47-F, StemLib-48-F, StemLib-49-F, and StemLib-52-F (table S2) were mixed at equal molar ratio and were used as the forward primer, whereas primer StemLib-WT-R (table S2) was used as the reverse primer. In the second PCR, primers StemLib-WT-F and StemLib-111-R were used. The products of the first and second PCRs were mixed at a molar ratio of 7:1 (the product of the first PCR to the product of the second PCR). This mixture was the insert for the single mutant library.

Insert for the double mutant library

The double mutant library was generated by two PCRs using the WT plasmid as template. In the first PCR, primers StemLib-42-F, StemLib-45-F, StemLib-46-F, StemLib-47-F, StemLib-48-F, StemLib-49-F, and StemLib-52-F (table S2) were mixed at equal molar ratio and were used as the forward primer, whereas primer StemLib-111-R (table S2) was used as the reverse primer. In the second PCR, primers StemLib-42/45-F, StemLib-42/46-F, StemLib-42/47-F, StemLib-42/48-F, StemLib-42/49-F, StemLib-42/52-F, StemLib-45/46-F, StemLib-45/47-F, StemLib-45/48-F, StemLib-45/49-F, StemLib-45/52-F, StemLib-46/47-F, StemLib-46/48-F, StemLib-46/49-F, StemLib-46/52-F, StemLib-47/48-F, StemLib-47/49-F, StemLib-47/52-F, StemLib-48/49-F, StemLib-48/52-F, and StemLib-49/52-F (table S2) were mixed at equal molar ratio and were used as the forward primer, whereas primer StemLib-WT-R (table S2) was used as the reverse primer. The products of the first and second PCRs were mixed at a molar ratio of 1:3 (the product of the first PCR to the product of the second PCR). This mixture was the insert for the double mutant library.

Vector generation

The vector for the mutant libraries was created by PCR using the WT plasmid as template and primers StemLib-VF and StemLib-VR (table S2).

Restriction digestion, ligation, and transformation

Both the vector and inserts were digested with BsmBI (New England Biolabs). Ligation was performed for each mutant library using T4 DNA ligase (New England Biolabs). The ligated products were transformed into MegaX DH10B T1R Electrocomp cells (Thermo Fisher Scientific). At least one million colonies were collected for each mutant library. Plasmid mutant libraries were purified from the bacteria colonies using Maxiprep Plasmid Purification (Clontech Laboratories).

**HA deep mutational scanning of H3/HK68**

Virus mutant libraries were rescued from the plasmid mutant libraries by transfecting HEK293T/MDCK-SIAT1 cells co-culture (ratio of 6:1) using lipofactamine 2000 (Thermo Fisher Scientific) according to the manufacturer's instructions. Transfection for the single mutant library was performed in a T75 $cm^2$ flask, and in a T225 $cm^2$ flask for the double mutant libraries. Two independent transfections were performed for each mutant library. Subsequently, four virus mutant libraries were produced, namely replicate 1 and 2 of the single mutant library, and replicate 1 and 2 of the double mutant library. For passaging of each virus mutant library, a monolayer MDCK-SIAT1 cells in a T75 $cm^2$ flask (for the single mutant library) or in a T225 $cm^2$ flask (for the double mutant library) were infected with an MOI of 0.05. At 2 hours post-infection, infected cells were washed three times with PBS followed by the addition of fresh medium. The virus mutant library was harvested at 24 hours post-infection. The harvested virus mutant library was also the post-selection library in this study. For profiling antibody resistance, antibody was added to the media at the indicated concentration throughout the course of infection. The antibody was incubated with the virus mutant library for 1 at room temperature, before the mixture of antibody-virus mutant library being added to the cells. Each virus mutant library (replicate 1 and 2 of the single mutant library, and replicate 1 and 2 of the double mutant library) was passaged in 5 conditions, namely no antibody, 2 μg $mL^{-1}$ of CR9114 IgG, 10 μg $mL^{-1}$ of CR9114 IgG, 0.3 μg $mL^{-1}$ of FI6v3 IgG and 2.5 μg $mL^{-1}$ of FI6v3 IgG.

**HA sequencing library preparation for H3/HK68**

Viral RNA was extracted using QIAamp Viral RNA Mini Kit (QIAGEN). The extracted RNA was then reverse transcribed to cDNA using Superscript III reverse transcriptase (Thermo Fisher Scientific). The plasmid mutant libraries or the cDNA from the post-infection viral mutant libraries were amplified by PCR using primers: 5'-CAC TCT TTC CCT ACA CGA CGC TCT TCC GAT CTA CAA GCA GCA GAT CTT AAA AGC-3' and 5'-GAC TGG AGT TCA GAC GTG

TGC TCT TCC GAT CTT CTC AAA CAG CTT GTT CAT TTC-3'. A second PCR was performed to add the rest of the adaptor sequence and index to the amplicon using primers: 5'-AAT GAT ACG GCG ACC ACC GAG ATC TAC ACT CTT TCC CTA CAC GAC GCT-3’ and 5'-CAA GCA GAA GAC GGC ATA CGA GAT XXX XXX GTG ACT GGA GTT CAG ACG TGT GCT-3'. Positions annotated by an “X” represented the nucleotides for the index sequence:

Single mutant library (Plasmid): 5'-GTA GCC-3'

Double mutant library (Plasmid): 5'-TAC AAG-3'

Single mutant library (no antibody, replicate 1): 5'-TTG ACT-3'

Single mutant library (no antibody, replicate 2): 5'-GGA ACT-3'

Double mutant library (no antibody, replicate 1): 5'-TGA CAT-3'

Double mutant library (no antibody, replicate 2): 5'-GGA CGG-3'

Single mutant library (2 μg $mL^{-1}$ of CR9114 IgG, replicate 1): 5'-CTC TAC-3'

Single mutant library (2 μg $mL^{-1}$ of CR9114 IgG, replicate 2): 5'-GCG GAC-3'

Double mutant library (2 μg $mL^{-1}$ of CR9114 IgG, replicate 1): 5'-TTT CAC-3'

Double mutant library (2 μg $mL^{-1}$ of CR9114 IgG, replicate 2): 5'-GGC CAC-3'

Single mutant library (10 μg $mL^{-1}$ of CR9114 IgG, replicate 1): 5'-CGA AAC-3'

Single mutant library (10 μg $mL^{-1}$ of CR9114 IgG, replicate 2): 5'-CGT ACG-3'

Double mutant library (10 μg $mL^{-1}$ of CR9114 IgG, replicate 1): 5'-CCA CTC-3'

Double mutant library (10 μg $mL^{-1}$ of CR9114 IgG, replicate 2): 5'-GCT ACC-3'

Single mutant library (0.3 μg $mL^{-1}$ of FI6v3 IgG, replicate 1): 5'-GTA GCC-3'

Single mutant library (0.3 μg $mL^{-1}$ of FI6v3 IgG, replicate 2): 5'-TAC AAG-3'

Double mutant library (0.3 μg $mL^{-1}$ of FI6v3 IgG, replicate 1): 5'-TTG ACT-3'

Double mutant library (0.3 μg $mL^{-1}$ of FI6v3 IgG, replicate 2): 5'-GGA ACT-3'

Single mutant library (2.5 μg $mL^{-1}$ of FI6v3 IgG, replicate 1): 5'-TGA CAT-3'

Single mutant library (2.5 μg $mL^{-1}$ of FI6v3 IgG, replicate 2): 5'-GGA CGG-3'

Double mutant library (2.5 μg $mL^{-1}$ of FI6v3 IgG, replicate 1): 5'-CTC TAC-3'

Double mutant library (2.5 μg $mL^{-1}$ of FI6v3 IgG, replicate 2): 5'-GCG GAC-3'

All final PCR products, except those from FI6v3 selections, were mixed (14 samples total) and submitted for next-generation sequencing using one lane of Illumina MiSeq PE300. The final PCR products from FI6v3 selections were mixed (8 samples total) and submitted for next-generation sequencing using 10% of one lane of Illumina MiSeq PE300.

**HA deep mutational scanning of H1/SI06 and H1/Mich15**

The single mutant libraries of H1/SI06 (A/Solomon Islands/3/2006) and H1/Mich15 (A/Michigan/45/2015) were constructed in the same manner as the single mutant library of H3/HK68 (see above). The primers are listed in table S3. Virus mutant libraries of H1/SI06 and H1/Mich15 were rescued with all non-HA segments from H1/WSN. Deep mutational scanning was performed as described for the single mutant library of H3/HK68, except only two passage conditions were used, namely no antibody and 0.3 μg $mL^{-1}$ of FI6v3 IgG. Of note, H1 is more sensitive to FI6v3 as compared to H3, so lower concentrations were used in the deep mutational scanning of H1, otherwise the virus would not survive. HA sequencing library for H1/SI06 and H1/Mich15 was prepared as described for H3/HK68, except different primers were used for PCR of plasmid mutant libraries and cDNA. For H1/SI06, the plasmid mutant libraries or the cDNA from the post-infection viral mutant libraries were amplified by PCR using primers: 5'-CAC TCT TTC CCT ACA CGA CGC TCT TCC GAT CTC TAT GCT GCG GAC CAA AAA AGC-3' and 5'-GAC TGG AGT TCA GAC GTG TGC TCT TCC GAT CTT CTC ATA CAG ATT CTT CAC ATT-3'. For H1/Mich15, the plasmid mutant libraries or the cDNA from the post-infection viral mutant libraries were amplified by PCR using primers: 5'-CAC TCT TTC CCT ACA CGA CGC TCT TCC GAT CTA TAT GCA GCC GAC CTG AAG AGC-3' and 5'-GAC TGG AGT TCA GAC GTG TGC TCT TCC GAT CTT TTC ATA CAA GTT CTT CAC ATT-3'. The index sequence for the second PCR was as follows:

H1/SI06 mutant library (no antibody, replicate 1): 5'-CGT GAT-3'

H1/SI06 mutant library (no antibody, replicate 2): 5'-ACA TCG-3'

H1/Mich15 mutant library (no antibody, replicate 1): 5'-GCC TAA-3'

H1/Mich15 mutant library (no antibody, replicate 2): 5'-TGG TCA-3'

H1/SI06 mutant library (0.3 μg $mL^{-1}$ of FI6v3 IgG, replicate 1): 5'-CAC TGT-3'

H1/SI06 mutant library (0.3 μg $mL^{-1}$ of FI6v3 IgG, replicate 2): 5'-ATT GGC-3'

H1/Mich15 mutant library (0.3 μg $mL^{-1}$ of FI6v3 IgG, replicate 1): 5'-GAT CTG-3'

H1/Mich15 mutant library (0.3 μg $mL^{-1}$ of FI6v3 IgG, replicate 2): 5'-TCA AGT-3'

H1/SI06 mutant library (Plasmid): 5'-CTG ATC-3'

H1/Mich15 mutant library (Plasmid): 5'-AAG CTA-3'

The final PCR products from H1/SI06 and H1/Mich15 selections were mixed (10 samples total) and submitted for next-generation sequencing using 30% of one lane of Illumina MiSeq PE300.

**HA sequencing data analysis for H3/HK68, H1/SI06, and H1/Mich15**

For each paired-end read, the positions of randomized codon were first identified by the internal barcode. A paired-end read would be discarded if the corresponding forward and reverse reads did not match at the internal barcode positions or at the randomized codon. This procedure was not applied when analyzing the mutant libraries of H1/SI06 and H1/Mich15, because an internal barcode was not used in the construction of the mutant libraries of H1/SI06 and H1/Mich15. Each mutation was called by comparing individual paired-end reads to the WT reference sequence. Sequencing data for each library were processed independently. For a mutant *i* in mutant library *n* of sample *t* (in this study, n could be the single mutant library or double mutant library, and t could be input plasmid library or library that was selected under a specified condition):

Occurrence $\text{frequency}_{i,n,t}$ = (Read $\text{count}_{i,n,t}$ + 1)/$\text{Coverage}_{n,t}$, where Read $\text{count}_{i,n,t}$ represents the number of read in mutant library *n* of sample *t* that carried mutation *i* and $\text{coverage}_n$ represents the sequencing coverage of the mutant library *n* of sample *t*.

Similarly, Occurrence $\text{frequency}_{WT,n,t}$ = (Read $\text{count}_{WT,n,t}$ + 1)/$\text{Coverage}_{n,t}$, where Read $\text{count}_{WT,n}$ represents the number of read that matches with the WT sequence in mutant library *n* of sample *t* and $\text{coverage}_{n,t}$ represents the sequencing coverage of the mutant library *n* of sample *t*.

Subsequently, Relative $\text{frequency}_{i,n,t}$ = (Occurrence $\text{frequency}_{i,n,t}$)/(Occurrence $\text{frequency}_{WT,n,t}$), and Relative $\text{fitness}_{i,n}$ = (Relative $\text{frequency}_{i,n,post\text{-}selection}$)/(Relative $\text{frequency}_{i,n,plasmid}$). Of note, a pseudocount was added to the read count when computing the occurrence frequency to avoid division by zero during the calculations of relative frequency and of relative fitness. For each mutant library, a given mutant would be discarded if the number of reads in the plasmid mutant library were <20 (<0.05% input frequency in the single mutant library and <0.003% input frequency in the double mutant library). The reported relative fitness was computed by averaging the relative fitness between replicates.

Relative resistance is computed by the ratio between the relative fitness with antibody selection and the relative fitness without antibody selection:

Relative $\text{resistance}_{i,n}$ = (Relative $\text{fitness}_{i,n,with\ antibody}$)/ (Relative $\text{fitness}_{i,n,without\ antibody}$)

Of note, relative resistance values could not be compared between CR9114 and FI6v3 selections since the strengths of selection pressures from CR9114 and FI6v3 are not normalized in this experiment.

**Deep mutational scanning of H1/WSN HA and H3/Perth09 HA**

The codon-mutant libraries of H1N1 A/WSN/1933 (H1/WSN) HA mutant virus libraries are those described in (*38*). The codon-mutant libraries of H3N2 A/Perth/16/2009 (H3/Perth09) HA mutant virus libraries are those described in (*39*). Details of the library generation and sequencing statistics of the mutant libraries can be found in (*38, 39*).

We performed antibody selections of the H1/WSN mutant virus libraries with 50 ng $mL^{-1}$, 70 ng $mL^{-1}$, and 100 ng $mL^{-1}$ of CR9114 IgG. We selected the H3/Perth09 mutant virus libraries with 5 µg $mL^{-1}$, 12 µg $mL^{-1}$, 13 µg $mL^{-1}$, and 15 µg $mL^{-1}$ of FI6v3 IgG. These antibody selection experiments were performed as described previously (*22, 31*). Briefly, we incubated $10^6$ $TCID_{50}$ per ml of mutant virus library with an equal volume of antibody at the intended concentration at 37°C for 1.5 hours. We also included a mock selection control wherein the mutant virus libraries were incubated with Influenza Growth Media (Opti-MEM supplemented with 0.01% heat-inactivated FBS, 0.3% BSA, 100 U $mL^{-1}$ of penicillin, and 100 µg $mL^{-1}$ of streptomycin). We used the H1/WSN mutant virus-antibody mixture to infect MDCK-SIAT1 cells, and the H3/Perth09 mutant virus-antibody mixture to infect MDCK-SIAT1-TMPRSS2 cells. The media was changed to fresh Influenza Growth Media (as above, except with 0.5% heat-inactivated FBS for the H1/WSN mutant viruses) at two hours post-infection. At 15 hours post-infection, we proceeded with library sequencing preparation.

We estimated the fraction of the mutant virus libraries surviving antibody selection (i.e. *fraction surviving*), as described previously (*22*). Briefly, we made duplicate 10-fold serial dilutions of the virus libraries to create a standard curve of infectivity. We then performed qPCR of the standard curve and antibody-selected samples using the NP and GAPDH primers provided in (*31*). We used linear regression to fit a line to relate the logarithm of the viral infectious dose from the standard curve to the difference in Ct values between NP and GAPDH. We then estimated the fraction of each viral library remaining infectious after antibody treatment using this relationship. The fraction of each library that remained infectious ranged from 0.14% to 2.3% for the H1/WSN libraries treated with FI6v3, 0.17% to 1% for the H1/WSN libraries treated with CR9114, and 4.3% to 7.4% for the H3/Perth09 libraries treated with FI6v3.

**HA sequencing library preparation for H1/WSN and H3/Perth09**

The libraries were prepared for deep sequencing as described in (*31, 39*). We extracted viral RNA using a Qiagen RNeasy Plus Mini Kit (QIAGEN). The extracted RNA was reverse-transcribed using AccuScript Reverse Transcriptase (Agilent) using H1/WSN HA- or Perth/2009 HA-specific primers, provided in (*31, 39*). Subsequent amplification of the cDNA and Illumina sequencing preparation were performed using a barcoded-subamplicon approach previously described (*28, 38*). All samples were submitted for next-generation sequencing on both lanes of an Illumina HiSeq 2500 using 2 x 250 bp paired-ends reads in rapid-run mode.

**HA sequencing data analysis for H1/WSN and H3/Perth09**

The dms_tools2 software package (*51*) (https://jbloomlab.github.io/dms_tools2/, version 2.3.0) was used to analyze the deep sequencing data.

**Construction of individual HA mutants**

Individual mutants for validation experiments were constructed using the QuikChange XL Mutagenesis kit (Stratagene) according to the manufacturer's instructions.

**Influenza microneutralization assay**

Microneutralization assay was performed as previously described (*26*). Briefly, two-fold serial dilutions of each antibody were incubated with 200 $TCID_{50}$ for 1 hour. MDCK-SIAT1 cells were washed twice with PBS and inoculated with virus-antibody mixtures. Cytopathic effect (CPE) was recorded at 72 hours post-inoculation. For the microneutralization assay of CR9114 and FI6v3 to H3/HK68, minimum inhibitory concentration (MIC) was the lowest concentration of the antibody that prevented observable CPE.

**Plaque morphology**

Confluent MDCK cells in 6-well plates were infected with 10-fold serial dilutions of virus in 1 ml infection medium for 1 hour. Cells were washed and overlaid with MEM with 0.3% BSA, 0.5% agarose, 1% Penicillin-Streptomycin and 1 µg/ml TPCK-trypsin and incubated at 37°C for 2 days. Cells were fixed with 4% formaldehyde overnight and the plaques were visualized by staining with 0.2% crystal violet solution.

**pH fusion assay**

The HA activation pH was determined by syncytium formation in Vero cells. Monolayers of Vero cells were infected with influenza viruses at a MOI of 10 PFU/cell for 1 hour. Six hours after infection, cells were incubated with infection medium containing 5 µg/mL TPCK- trypsin followed by pH adjusted PBS for 5 min at 37°C. Cells were overlaid with MEM containing 5% fetal bovine serum and further incubated at 37°C. When syncytium formation was identified microscopically, the cells were fixed and stained with the Differential Quick Stain Kit (EMS).

**Analysis of natural amino acid variants**

HA protein sequences were downloaded from Influenza Research Database (www.fludb.org/) (*33*). Sequence alignment was performed by MAFFT version 7.157b (*52*). HA protein sequences from human H3N2 isolates that were sequenced without any passaging were downloaded from Global Initiative for Sharing Avian Influenza Data (GISAID; http://gisaid.org). Passaging history was determined by parsing regular expression in the FASTA header as described previously (*34, 53*). Sequence logos were generated by WebLogo (http://weblogo.berkeley.edu/logo.cgi) (*54*). A total of 4625 human H3N2 HA sequences, 81 human H2N2 HA sequences, and 65 avian H2N2 HA sequences were included in our analysis. For the analysis of human H3N2 sequences, occurrence frequency of each amino acid variant $i$ at residue $n$ was computed by:

$$occurrence\ frequency_{i,n} = \sum_{t} \frac{occurrence\ frequency_{i,n,t}}{Total\ number\ of\ years}$$

where occurrence frequency$_{i,n,t}$ represents the occurrence frequency of amino acid variant *i* at residue *n* in year *t*, and total number of years represents the number of years being sampled. In this case, the total number of years equals 48 (from 1968 to 2015). Calculating occurrence frequency in this manner will avoid temporal sampling bias.

**Recombinant HA expression and purification**

H3/HK68 HAs (WT or mutants) were prepared for binding experiments as previously described (*41*). Briefly, the ectodomain of HA, which corresponds to 11–329 (HA1) and 1–176 (HA2) based on H3 numbering was fused with an N-terminal gp67 signal peptide and a C-terminal tag consisting of (from N- to C-terminus) BirA biotinylation site, thrombin cleavage site, foldon trimerization domain, and $His_6$ tag, and cloned into a customized baculovirus transfer vector (*41*). Recombinant bacmid DNA was generated using the Bac-to-Bac system (Thermo Fisher Scientific). Baculovirus was generated by transfecting purified bacmid DNA into Sf9 cells using FuGene HD (Promega). H1/WSN HA was expressed by infecting suspension cultures of High Five cells with baculovirus at an MOI of 5 to 10 and incubating at 28°C shaking at 110 rpm for 72 hours. The supernatant was concentrated. HA0 was purified by Ni-NTA and buffer exchanged into 20 mM Tris-HCl pH 8 and 150 mM NaCl. For binding experiments, HA0 was biotinylated as described (*55*) and purified by size exclusion chromatography on a Hiload 16/90 Superdex 200 column (GE Healthcare) in 20 mM Tris pH 8.0, 150 mM NaCl, and 0.02% NaN3. For crystallization, the HA0 was treated with trypsin (New England Biolabs) to remove the C-terminal tag (BirA biotinylation site, thrombin cleavage site, trimerization domain, and $His_6$ tag) and to produce the cleaved mature HA (HA1/HA2). The trypsin-digested HA was then purified by size exclusion chromatography on a Hiload 16/90 Superdex 200 column (GE Healthcare) in

20 mM Tris pH 8.0, 150 mM NaCl, and 0.02% $NaN_3$ and concentrated to 9 mg/mL in 10 mM Tris pH 8.0, 50 mM NaCl, and 0.02% $NaN_3$.

**Biolayer interferometry binding assay**

Binding assay was performed by biolayer interferometry (BLI) using an Octet Red instrument (FortéBio) as described previously (*56*). Briefly, biotinylated HA0 at 50 μg/mL in 1x kinetics buffer (1x PBS, pH 7.4, 0.01% BSA and 0.002% Tween 20) was loaded onto streptavidin biosensors and incubated with the indicated concentration of Fab or IgG. The assay consisted of five steps: 1) baseline: 60 s with 1x kinetics buffer; 2) loading: 120 s with biotinylated HA0; 3) baseline: 60 s with 1x kinetics buffer; 4) association: 120 s with samples (Fab or IgG); and 5) dissociation: 120 s with 1x kinetics buffer. For estimating the exact $K_d$, a 1:1 binding model was used. In cases where the binding affinity was relatively weak ($K_d$ > 100 nM), a 1:1 binding model did not fit well due to the contribution of non-specific binding to the response curve. Subsequently, a 2:1 heterogeneous ligand model was used to improve the fitting.

**Crystallization and structural determination**

All H3/HK68 HA mutants were crystallized using the sitting drop vapor diffusion method with 500 μL reservoir solution containing 0.1 M sodium cacodylate pH 6.5, 5% PEG 8000, and 38% 2-methyl-2,4-pentanediol. Drops consisting 0.8 μL protein at 10 mg/ml + 0.8 μL precipitant were set up at 20 °C and crystals appeared within 3 days. This crystallization condition was as previously described for H3/HK68 receptor-binding site mutants (*30*). Diffraction data were collected at the Stanford Synchrotron Radiation Lightsource beamline 12-2 and indexed, integrated and scaled using HKL2000 (HKL Research) (*57*). The structure was solved by molecular replacement using Phaser (*58*) with PDB: 4FNK (*55*) as the molecular replacement model, then modeled using Coot (*59*) and refined using Refmac5 (*60*). Ramachandran statistics were calculated using MolProbity (*61*). For data collection and refinement statisitics, see table S1.

**Buried surface area calculation**

Solvent accessibility was computed by DSSP (*62*). Buried surface area (BSA) was calculated by subtracting the solvent accessibility of the apo form from that of the bound form. BSA was then normalized to the empirical scale reported in (*63*) to obtain the relative solvent accessibility (RSA).

***In vivo* pathogenesis and protection study**

Male and female 8-to-12-week-old C57BL/6J mice were obtained from The Scripps Research Institute rodent breeding colony and maintained in biosafety containment under pathogen-free conditions. All mouse experiments were approved by The Scripps Research Institute IACUC and complied with recommendations in the Guide for the Care and Use of Laboratory Animals of the National Research Council. All recombinant H3/HK68 (7:1 on H1/PR8 backbone) viruses were grown in MDCK cultures (approximately 40,000,000 cells per virus) for 72 hours in 1x MEM (Invitrogen) supplemented with 0.1% BSA, 1x Penicillin-Streptomycin, 1x L-glutamine (Invitrogen), and 1 μg $ml^{-1}$ (final) TPCK-treated trypsin (Sigma). Following incubation, supernatant was collected. Cell debris was pelleted by low-speed centrifugation (500 rcf for 5 min), and viruses isolated from the supernatant by high-speed centrifugation at 60,000 rcf. for 120 min. Virus pellets were resuspended in sterile 1x PBS containing 5% glycerol, aliquoted, and stored at -80°C. $TCID_{50}$ titers of all virus stocks were determined in MDCK cell cultures prior to infection studies.

For pathogenesis experiments, groups of n = 4 or 5 C57BL/6J mice under deep isoflurane anesthesia were inoculated intranasally with 25 μl doses of H3/HK68 (7:1 on H1/PR8 backbone)

WT or mutant viruses, or PBS, ranging from $10^4$ to $10^1$ $TCID_{50}$ $ml^{-1}$. Following infection, mice were monitored for body weight loss or clinical signs of infection every 24 hours up to 15 days post infection. Mice that reached 30% loss of original body weight on day 0, or were observed to have a severe/morbid clinical score on three consecutive days, were humanely euthanized. Resulting data were used to compare infectivity of WT and mutant viral stocks and calculate respective $mLD_{50}$ titers.

For prophylactic protection experiments, groups of n = 8 (4 male and 4 female) C57BL/6J mice were intravenously immunized with 100 µl of CR9114 at 10, 4, or 1 mg $kg^{-1}$ doses, or PBS control, via tail vein injection prior to challenge with a lethal dose of WT or respective mutant H3/HK68 viruses (7:1 on H1/PR8 backbone). 24 hours after immunization, all mice were infected intranasally, exactly as described above, with a dose equivalent to 25 $mLD_{50}$ of respective virus stocks. All mice were monitored for body weight loss or clinical signs of infection every 24 hours up to 15 days post infection. Mice that reached 30% loss of original body weight on day 0, or were observed to have a severe/morbid clinical score on three consecutive days, were humanely euthanized. Effectiveness of antibody protection was analyzed via Kaplan-Meier survival curves and paired analysis of each treatment group, relative to control, was conducted using Log-rank (Mantel-Cox) tests.

**Construction of yeast surface display library**

The insect cell expression plasmid that encodes WT CR9114 Fab (*26*) was used as the template to generate the CR9114 Fab yeast display construct. Specifically, the heavy chain and light chain of CR9114 were cloned into the dual promoter yeast expression plasmid that was described previously (*56*). To construct the insert for the CR9114 mutant library, a PCR was performed using the WT CR9114 Fab yeast expression plasmid as template. The insert was generated using an overlapping PCR strategy with two fragments. The first fragment of the insert was generated using primers: 5'-AAA AGT AGC GGA GGA ACA TCA AAC NNS TAC GCA ATC TCT TGG GTG CGG CA-3' and 5'-CTG AGC GTA GGC TGT ACT CCC MNN SNN SNN AGA GAT CCC GCC CAT CCA GTC-3'. The second fragment of the insert was generated using primers: 5'-GAC TGG ATG GGC GGG ATC TCT NNS NNS NNK GGG AGT ACA GCC TAC GCT CAG-3' and 5'-GAC ATC CAT CCC AGA GTA GTA ATA SNN GTT TCC GTG GCG AGC ACA AAA GT-3'. The two fragments were mixed at equal molar ratio and were used as the template for PCR using primers: 5'-AGC CCG GCA GTA GTG TCA AAG TCA GTT GTA AAA GTA GCG GAG GAA CAT CA-3' and 5'-GAG ACA GTG ACG GTT GTT CCC TGC CCC CAG ACA TCC ATC CCA GAG TAG TA-3'. The product of this overlapping PCR is the insert for the CR9114 mutant library. The vector for the CR9114 Fab mutant library was generated by PCR using the WT CR9114 Fab yeast expression plasmid as template and primers: 5'-TAC TAC TCT GGG ATG GAT GTC TGG GGG CAG GGA ACA ACC GTC ACT GTC TC-3' and 5'-TGA TGT TCC TCC GCT ACT TTT ACA ACT GAC TTT GAC ACT ACT GCC GGG CT-3'. The insert and vector were mixed and transformed into yeast strain EBY100 using a high-efficiency protocol as previously described (*64*). At least 50 million yeast colonies were collected to generate the CR9114 Fab yeast surface display library, which was resuspended in SDCAA (2.0% glucose, 0.67% yeast nitrogen base, 0.5% casamino acids, 0.54% disodium phosphate, 0.86% monosodium phosphate) with 15% glycerol and stored at −80 °C until used.

**Selection of yeast surface display library**

Selection was performed as previously described (*56*). Briefly, for each round of selection, $10^9$ yeast cells from the frozen stock were cultured in 250 mL SDCAA for 16 hours at 28 °C with shaking at 250 rpm. Yeast cells were then pelleted and resuspended in 100 ml SGR-CAA (20 g $L^{-1}$ galactose, 20 g $L^{-1}$ raffinose, 1 g $L^{-1}$ dextrose, 6.7 g $L^{-1}$ yeast nitrogen base, 5 g $L^{-1}$ casamino acids, 5.4 g $L^{-1}$ $Na_2HPO_4$ and 8.56 g $L^{-1}$ $Na_2HPO_4$). Yeast cells were cultured for 24 hours at 18 °C with shaking at 250 rpm to reach an $OD_{600}$ of 1.2 to 1.6. 8 mL of the yeast culture was spun down, washed twice with PBS, and resuspended in 5 ml PBS. Biotinylated trimeric HA was incubated with streptavidin-PE (eBioscience, San Diego, CA) at a molar ratio of 1:4 for 15 min. Of note, Streptavidin PE was buffer exchanged into PBS before use to remove $NaN_3$, which is toxic to the yeast cells. The biotinylated trimeric HA-strepavidin-PE complex was added to the yeast cells in PBS with a final concentration of 10 nM, or 0.4 nM for round 2 of WT H3/HK68 selection, or 0.016 nM for round 3 of WT H3/HK68 selection. After incubating at 4 °C overnight with head-to-head rotation, the yeast cells were pelleted, washed with PBS, resuspended in 5 mL PBS, and subjected to fluorescence-activated cell sorting (FACS) at TSRI Flow Cytometry Core Facility. The sorted yeast cells were recovered by plating on the SDCAA agar plates. Yeast colonies were collected after 2 days of incubation at 30 °C, resuspended in YPD with 15% glycerol, and stored at −80 °C until used.

**Next-generation sequencing of yeast surface display library**

Plasmid was extracted from at least $10^7$ yeast cells per sample using Zymoprep Yeast Plasmid Miniprep II (Zymo Research) according to the manufacturer's instructions. The mutated region on the CR9114 heavy chain was amplified by PCR using primers: 5'-CAC TCT TTC CCT ACA CGA CGC TCT TCC GAT CTG TAA AAG TAG CGG AGG AAC ATC-3' and 5'-GAC TGG AGT TCA GAC GTG TGC TCT TCC GAT CTC AGA CAT CCA TCC CAG AGT AGT-3'. A second PCR was performed to add the rest of the adaptor sequence and index to the amplicon using primers: 5'-AAT GAT ACG GCG ACC ACC GAG ATC TAC ACT CTT TCC CTA CAC GAC GCT-3' and 5'-CAA GCA GAA GAC GGC ATA CGA GAT XXX XXX GTG ACT GGA GTT CAG ACG TGT GCT-3'. Positions annotated by an "X" represented the nucleotides for the index sequence:

Mutant library DNA insert: 5'-CGT GAT-3'

Yeast pre-selected mutant library: 5'-ACA TCG-3'

WT round 1: 5'-GCC TAA-3'

WT round 2: 5'-TGG TCA-3'

WT round 3: 5'-CAC TGT-3'

I45T round 1: 5’-CTG ATC-3'

I45T round 2: 5'-AAG CTA-3'

I45T round 3: 5'-GTA GCC-3'

I45M round 1: 5'-ATT GCC-3'

I45M round 2: 5'-GAT CTG-3'

I45M round 3: 5'-TCA AGT-3'

I45F round 1: 5'-TAC AAG-3'

I45F round 2: 5'-TTG ACT-3'

I45F round 3: 5'-GGA ACT-3'

For each paired-end read, the nucleotide sequence corresponding to the randomized region was extracted. If the nucleotide sequence at the randomized region was inconsistent between forward and reverse reads, the paired-end read would be discarded. In other words, at the randomized region, the reverse-complement of forward read must perfectly match the reverse read. The nucleotide sequence was translated into the amino-acid sequence. The occurrence of each amino-acid sequence was counted, with each paired-end read as one count.

**Code availability**

Computational scripts for analyzing the H3/HK68 deep mutational scanning data have been deposited to https://github.com/wchnicholas/HAstemEscape. Computational scripts for analyzing the H3/Perth09 and H1/WSN deep mutational scanning data have been deposited to https://github.com/jbloomlab/HA_stalkbnAb_MAP. Computational scripts for analyzing the CR9114 deep mutational scanning data have been deposited to https://github.com/wchnicholas/CR9114mut.

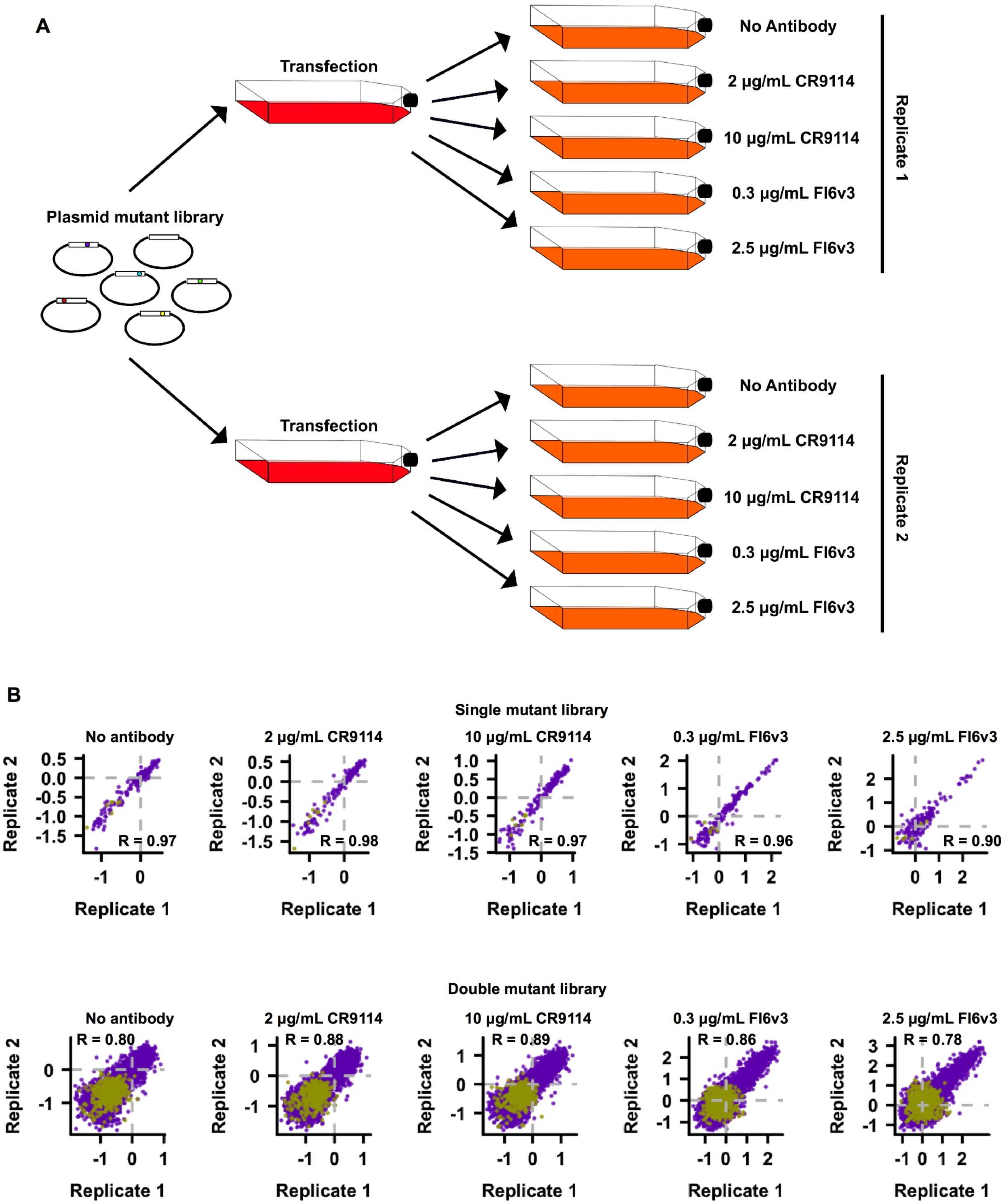


**Fig. S1. Experimental design and correlation between replicates. (A)** Schematic representation of the experimental design is shown. **(B)** Relative fitness of each mutant under different growth conditions is shown. Each data point represents one mutant. Pearson correlations (R) of the relative fitness of individual mutants between replicates are shown. Missense variants are colored in purple. Nonsense variants are colored in khaki green. The relative fitness for each mutant correlated well between biological replicates (Pearson correlation = 0.78 to 0.98), thereby demonstrating the high reproducibility of our deep mutational scanning.

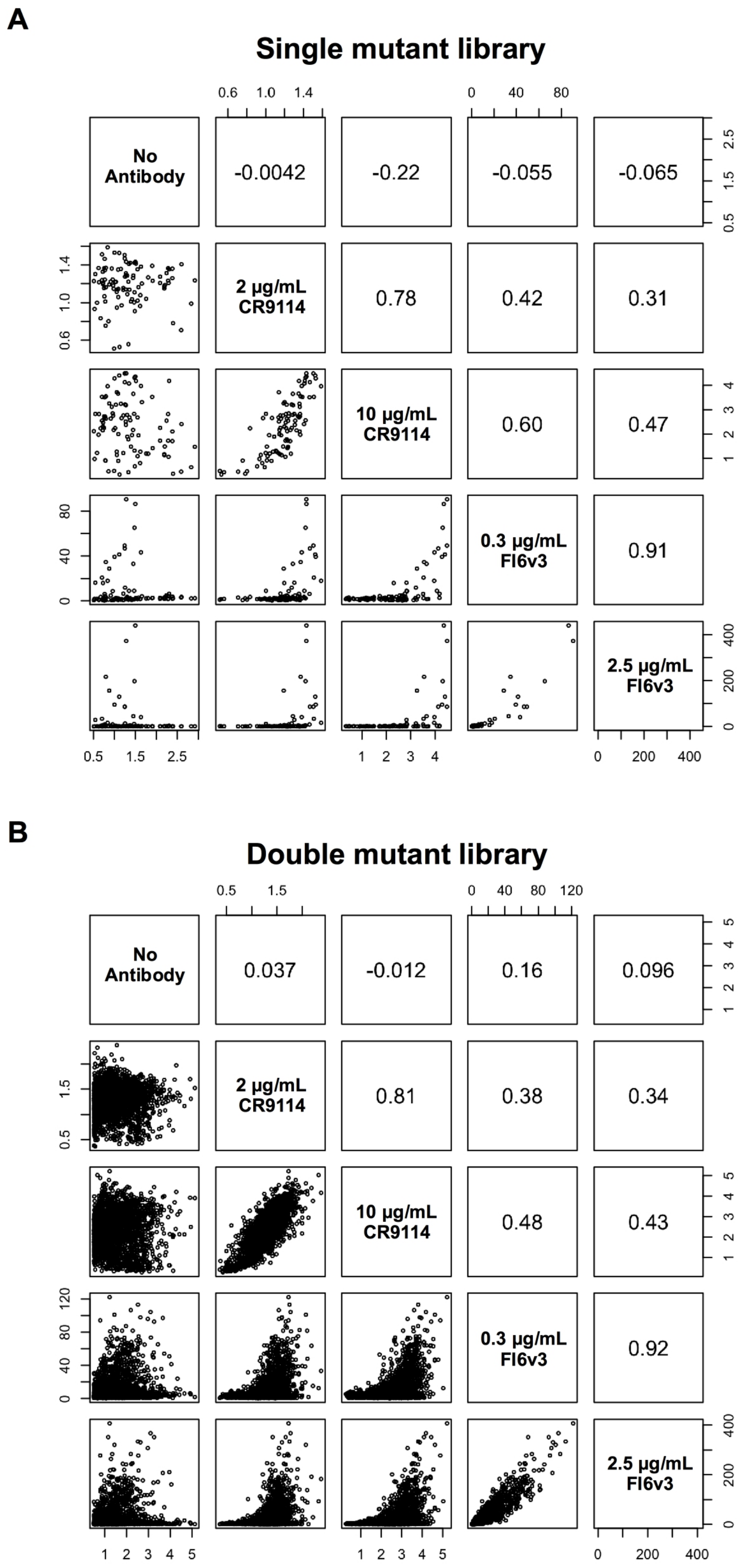


**Fig. S2. Relationships among relative fitness and relative resistance.** The relationships among relative fitness (no antibody), and relative resistance to 2 μg/mL CR9114 antibody, 10 μg/mL CR9114 antibody, 0.3 μg/mL FI6v3 antibody, and 2.5 μg/mL FI6v3 antibody for **(A)** single HA mutant virus library, and **(B)** double HA mutant virus library are shown as scatter plots. Each data point in the scatter plots represents a mutant. The Pearson correlation coefficients are indicated. The overall relative resistance profiles between CR9114 and FI6v3 were moderately correlated, as expected given that they target a broadly similar region of the HA.

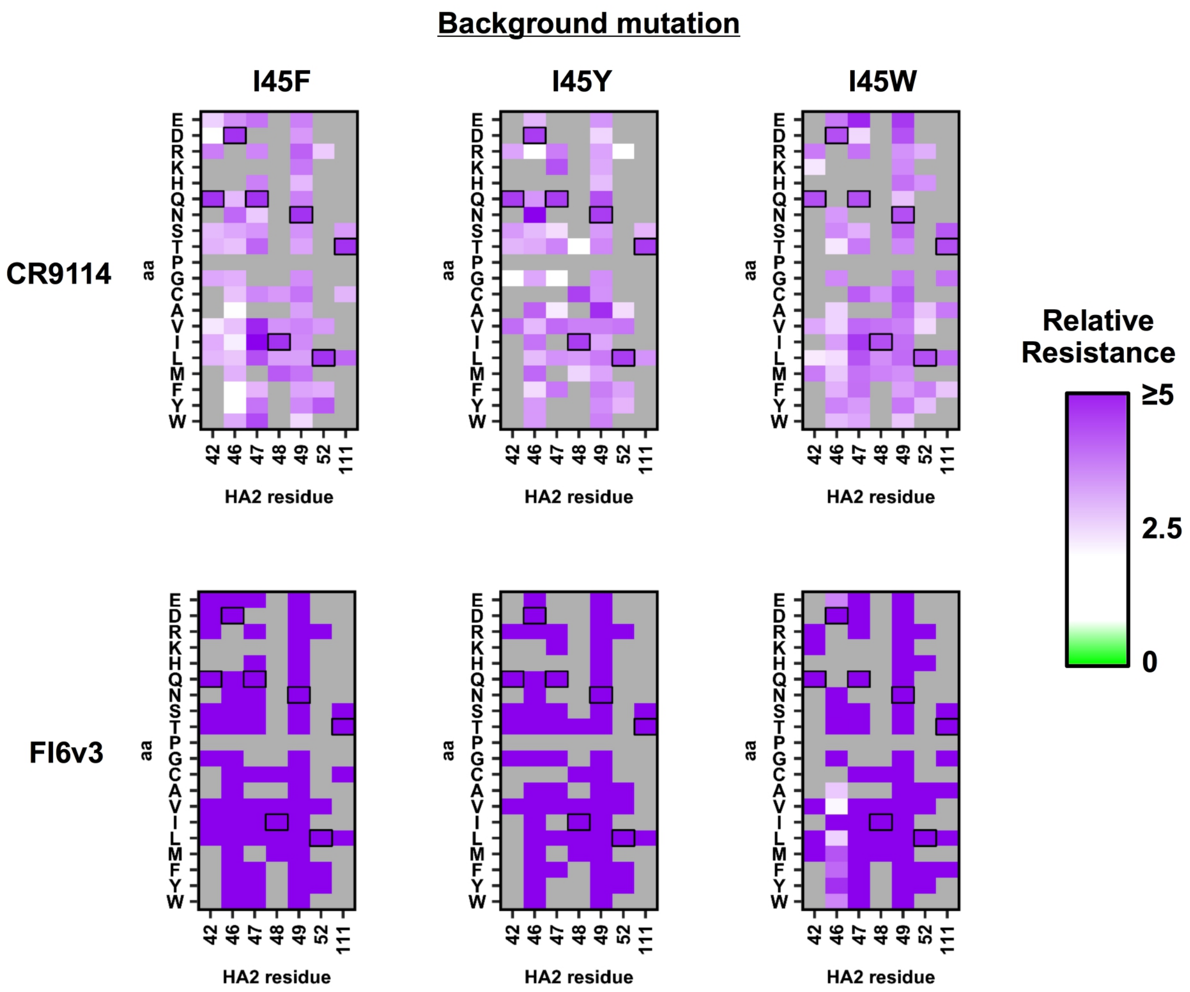


**Fig. S3. Relative resistance of selected sets of double mutants on H3/HK68 virus to HA stem antibodies.** Relative resistance of double mutants against 10 μg/mL CR9114 antibody or 2.5 μg/mL FI6v3 antibody is shown. Each data point in the heatmap represents a double mutant that is composed of the background mutation (as indicated at the top) plus a mutation (as indicated by the x/y-axes). For example, the top left data point of the top left heatmap represents the relative resistance of the I45F/Q42E double mutant against 10 μg/mL CR9114 antibody. Relative resistance for WT is set as 1. Virus mutants with a relative fitness of less than 0.5 are excluded from this analysis and are shown as grey. Residues correspond to WT sequence are boxed. When double mutants were composed of at least one strong resistance mutation (e.g. I45F, I45Y, and I45W), they showed strong resistance even if the other mutation did not confer resistance by itself. For example, while most single mutants at residues 47 and 49 did not have high relative resistance (Fig. 2C), mutations at residues 47 or 49 in combination with a mutation with high relative resistance, such as I45Y/F/W, resulted in high relative resistance of the virus to stem bnAbs.

**A**

**Human H3N2 isolates that were sequenced without any passaging**

| Strain | Accession | 42 | 45 | 46 | 47 | 48 | 49 | 52 | 111 |
|---|---|---|---|---|---|---|---|---|---|
| A/Virginia/66/2016 | EPI_ISL_241581 | Q | T | D | Q | I | N | L | T |
| A/Hawaii/105/2016 | EPI_ISL_244700 | Q | T | D | Q | I | N | L | T |

**B**

**Representative strains from different subtypes**

| Strain | 42 | 45 | 46 | 47 | 48 | 49 | 52 | 111 |
|---|---|---|---|---|---|---|---|---|
| B/Phuket/3073/2013 (B-Yamagata) | Q | I | N | K | I | T | L | E |
| B/Brisbane/60/2008 (B-Victoria) | Q | I | N | K | I | T | L | E |
| A/Hong Kong/3239/2008 (H9) | Q | I | D | K | I | T | V | H |
| A/turkey/Ontario/6118/1968 (H8) | Q | I | D | K | I | T | V | H |
| A/duck/Alberta/60/1976 (H12) | Q | I | D | N | M | Q | L | H |
| A/Taiwan/2/2013 (H6) | Q | I | D | G | I | T | V | H |
| A/California/04/2009 (H1) | Q | I | D | E | I | T | V | H |
| A/Japan/305/1957 (H2) | Q | F | D | G | I | T | V | H |
| A/Vietnam/1203/2004 (H5) | Q | I | D | G | V | T | V | H |
| A/Duck/England/56 (H11) | Q | I | D | Q | I | T | V | H |
| A/Gull/Maryland/704/77 (H13) | Q | I | D | Q | I | T | I | H |
| A/black-headed gull/Sweden/2/99 (H16) | Q | I | N | E | I | T | I | H |
| A/Brisbane/10/2007 (H3) | Q | I | D | Q | I | N | L | T |
| A/Duck/Czechoslovakia/1956 (H4) | Q | I | D | Q | I | N | L | T |
| A/Mallard/Gurijev/263/82 (H14) | Q | I | D | Q | I | N | L | T |
| A/Jiangxi-Donghu/346/2013 (H10) | Q | I | D | Q | I | T | L | A |
| A/Shanghai/2/2013 (H7) | Q | I | D | Q | I | T | L | A |
| A/shearwater/Australia/2576/1979 (H15) | Q | I | D | Q | I | T | L | A |

**Fig. S4. Sequence alignment of representative influenza strains.** The amino acid variants at the HA2 residues of interest in **(A)** two human H3N2 isolates that were sequenced without any passaging and **(B)** representative strains from different HA subtypes and from influenza B virus are shown.

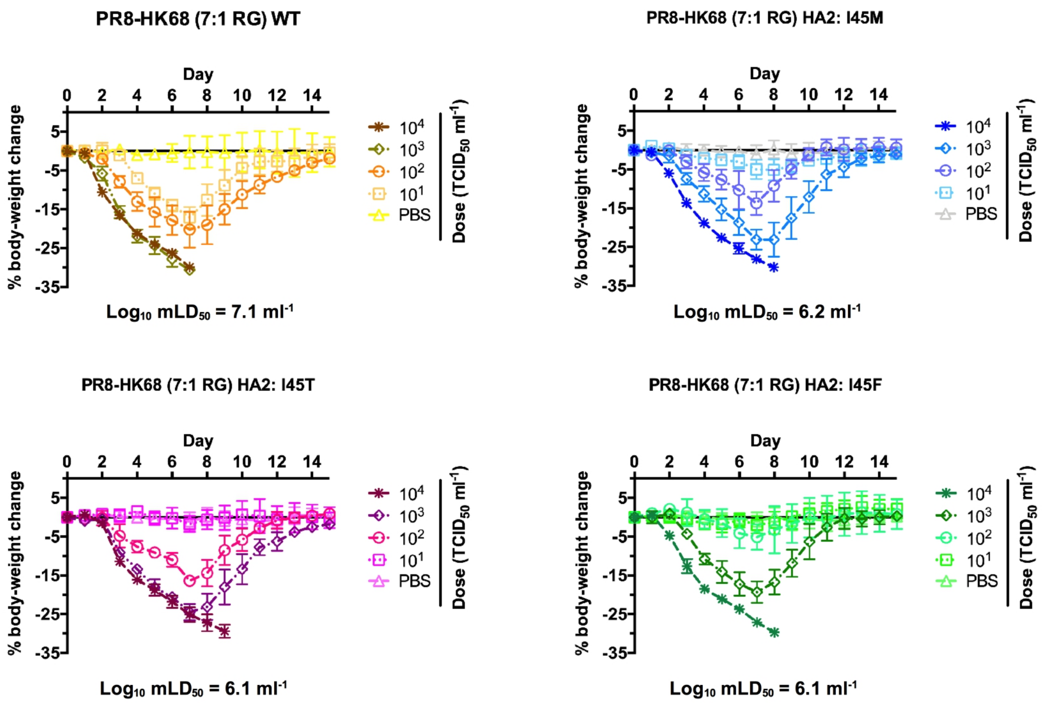


**Fig. S5. *In vivo* pathogenesis of antibody-resistant mutants as determined by weight loss in mice.** Mice were inoculated with the indicated dose ($TCID_{50}$ $ml^{-1}$) of the H3/HK68 (7:1 on H1/PR8 backbone) WT or mutant viruses. For each dose of WT or mutant viruses, a group of n = 4 or 5 C57BL/6J mice were used. Body-weight changes were measured. The mean is plotted and error bars represent standard deviation. The $mLD_{50}$ for each virus is indicated below the plots.

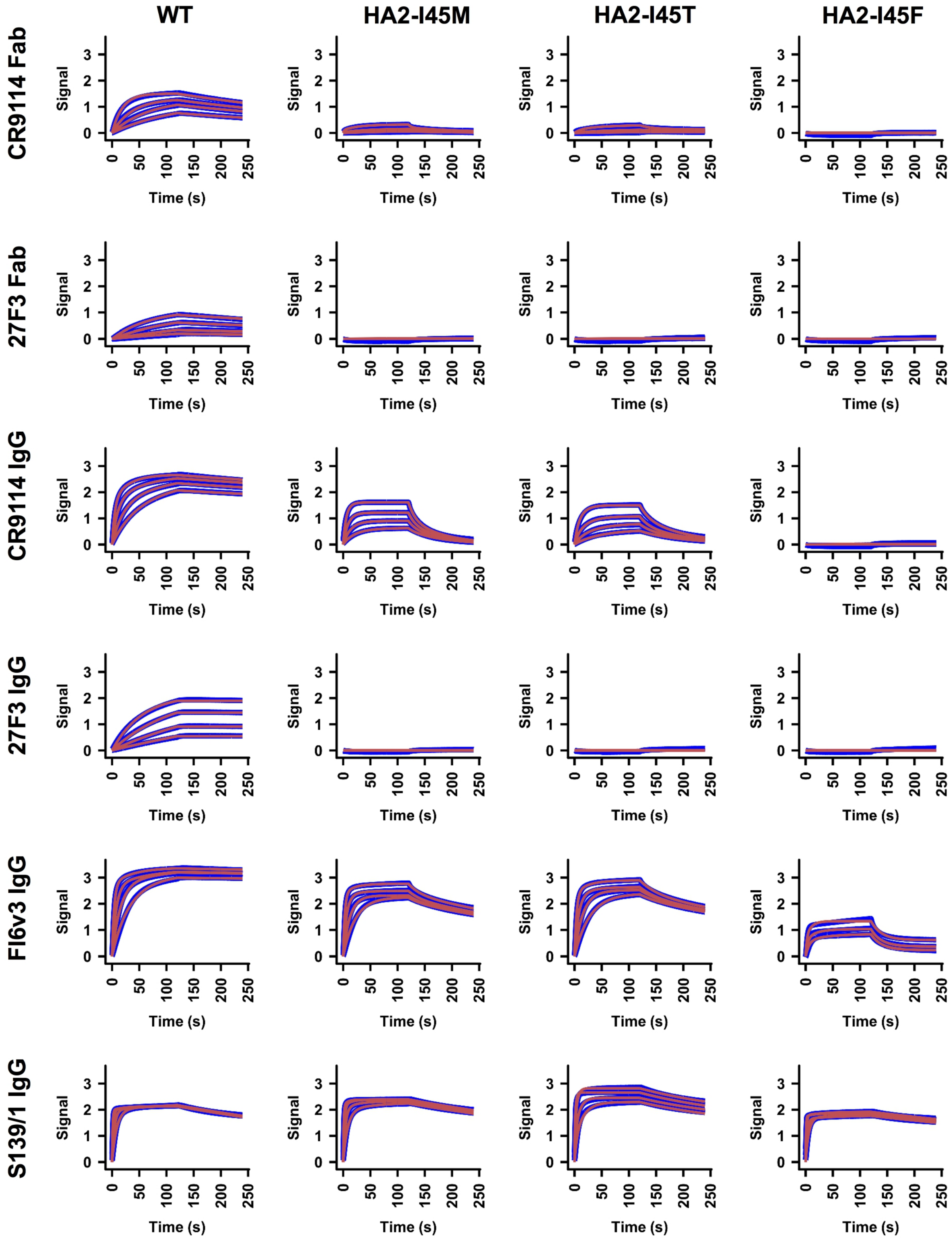


**Fig. S6. Sensorgrams for binding of Fabs and IgGs to HA2 mutants.** Binding kinetics of different Fabs and IgGs against recombinant H3/HK68 HA (WT or mutants) by biolayer interferometry (BLI). Y-axis represents the response. Blue lines represent the response curve and red lines represent the best fit model (1:1 binding model or 2:1 heterogeneous ligand model, see Methods). Binding kinetics were measured for three to four concentrations of Fab at 2-fold dilution ranging from 1,000 nM to 125 nM. Of note, the bivalency of IgG can significantly enhance its binding as compared to Fab. Such an observation has been described in our previous study of 27F3 (*35*).

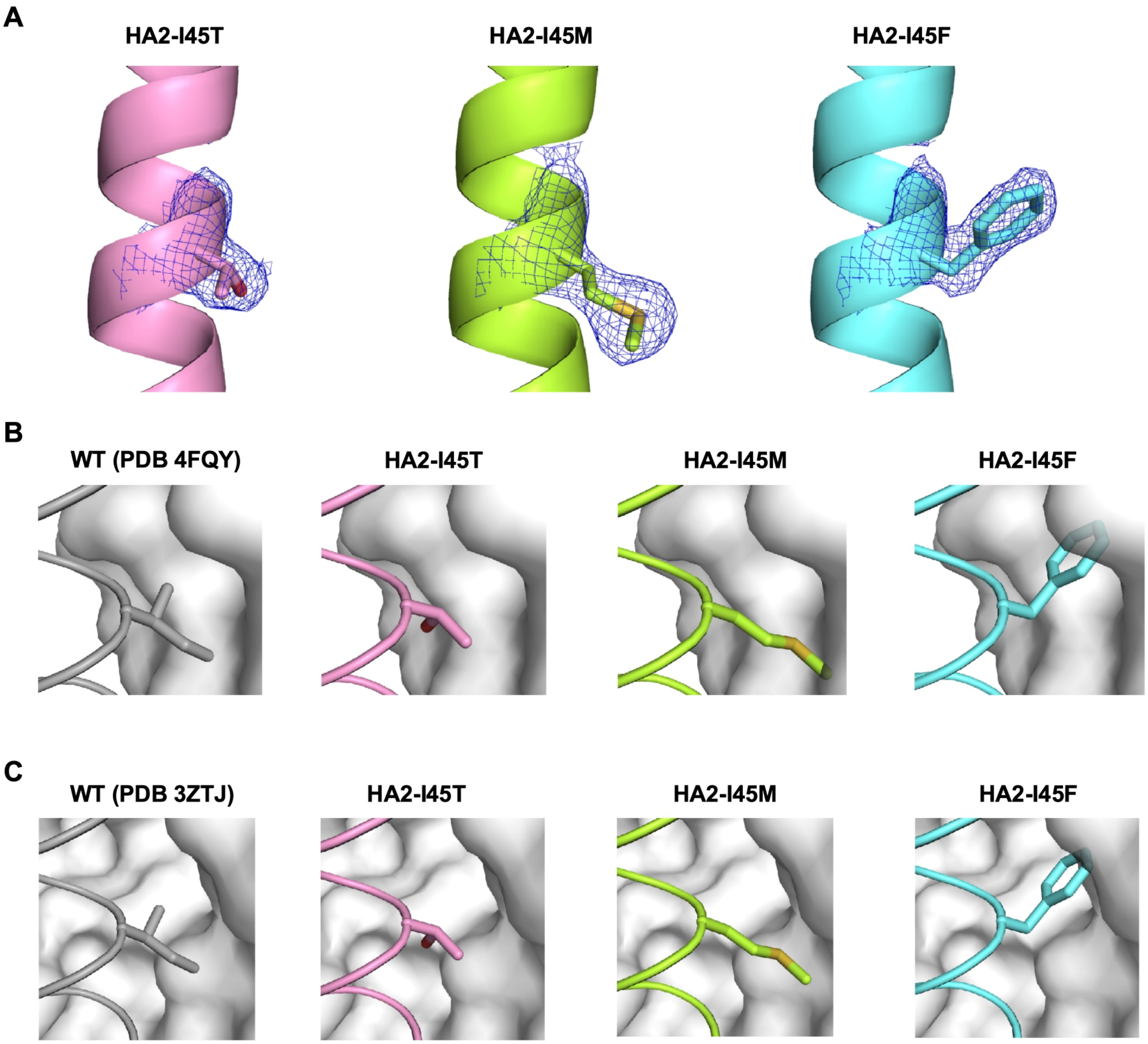


**Fig. S7. Structural characterization of H3/HK68 HA2 mutants. (A)** Final 2Fo-Fc electron density maps for HA2 residue 45 in different mutants are represented in a blue mesh and contoured at 1.0 σ. The side chains of HA2 residue 45 in different mutants are shown in stick representation. **(B)** To analyze the effect of different HA2 mutants on CR9114 binding, the apo structures of HA2 I45T, I45M, and I45F mutants were aligned with WT H3/HK68 HA in complex with CR9114 Fab (PDB 4FQY) (*26*). The backbone of helix A is shown in tube representation, side chain of residue 45 in stick representation, and CR9114 Fab in surface representation. **(C)** To analyze the effect of different HA2 mutants on FI6v3 binding, the apo structures of HA2 I45T,

I45M, and I45F mutants are aligned with the WT H3/HK68 HA in complex with FI6v3 Fab (PDB 3ZTJ) (*27*). The backbone of helix A is shown in tube representation, the side chain of residue 45 in stick representation, and FI6v3 Fab in surface representation.

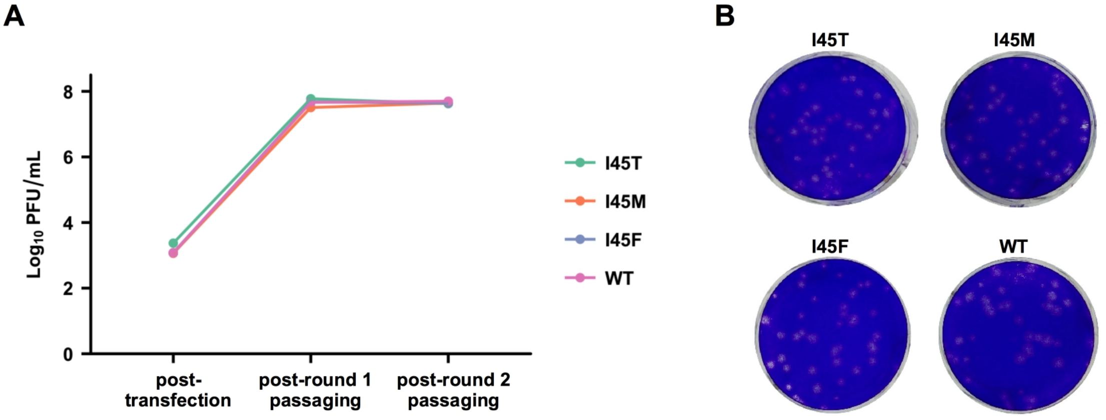


C

| Virus | Minimum inhibitory concentration (MIC) |
| --- | --- |
| HA2 I45T mutant | >400 μg/ml |
| HA2 I45M mutant | >400 μg/ml |
| HA2 I45F mutant | >400 μg/ml |
| WT | 5.9 μg/ml |

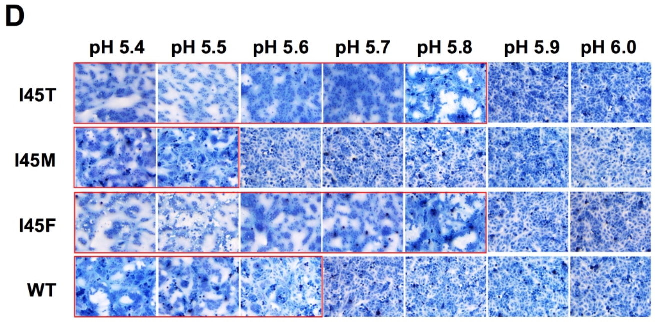


**Fig. S8. Characterization of A/Wuhan/359/95 (H3N2) HA2 mutants. (A)** Recombinant viruses were rescued at 293T cells and then passaged in MDCK cells at an MOI of 0.001 PFU/cell (for round 1 passaging) or 0.005 PFU/cell (for round 2 passaging). The titer for viral rescue (post-transfection) and post-passaging is shown. **(B)** Plaque morphologies of A/Wuhan/359/95 (H3/Wuhan95) HA2 mutants are shown. **(C)** The minimum inhibitory concentrations (MIC) of FI6v3 to H3/Wuhan95 HA2 mutants I45T, I45M, I45F, and wild type (WT) are shown. **(D)** The pH for HA activation was determined by syncytium formation in Vero cells after infection with recombinant H3/Wuhan95 viruses at an MOI of 10. Representative images (x10 magnification) of syncytium formation at indicated pH values are shown. The experiments were repeated twice with recombinant viruses after one passage in MDCK cells and once with recombinant viruses after two passages in MDCK cells. The fusion pHs of H3/Wuhan95 I45T, I45M, and I45F were pH 5.8, 5.5, and 5.8, respectively, which differed slightly from pH 5.6 of WT. Increase in fusion pH can contribute to resistance against HA stem bnAbs (*20, 65, 66*).

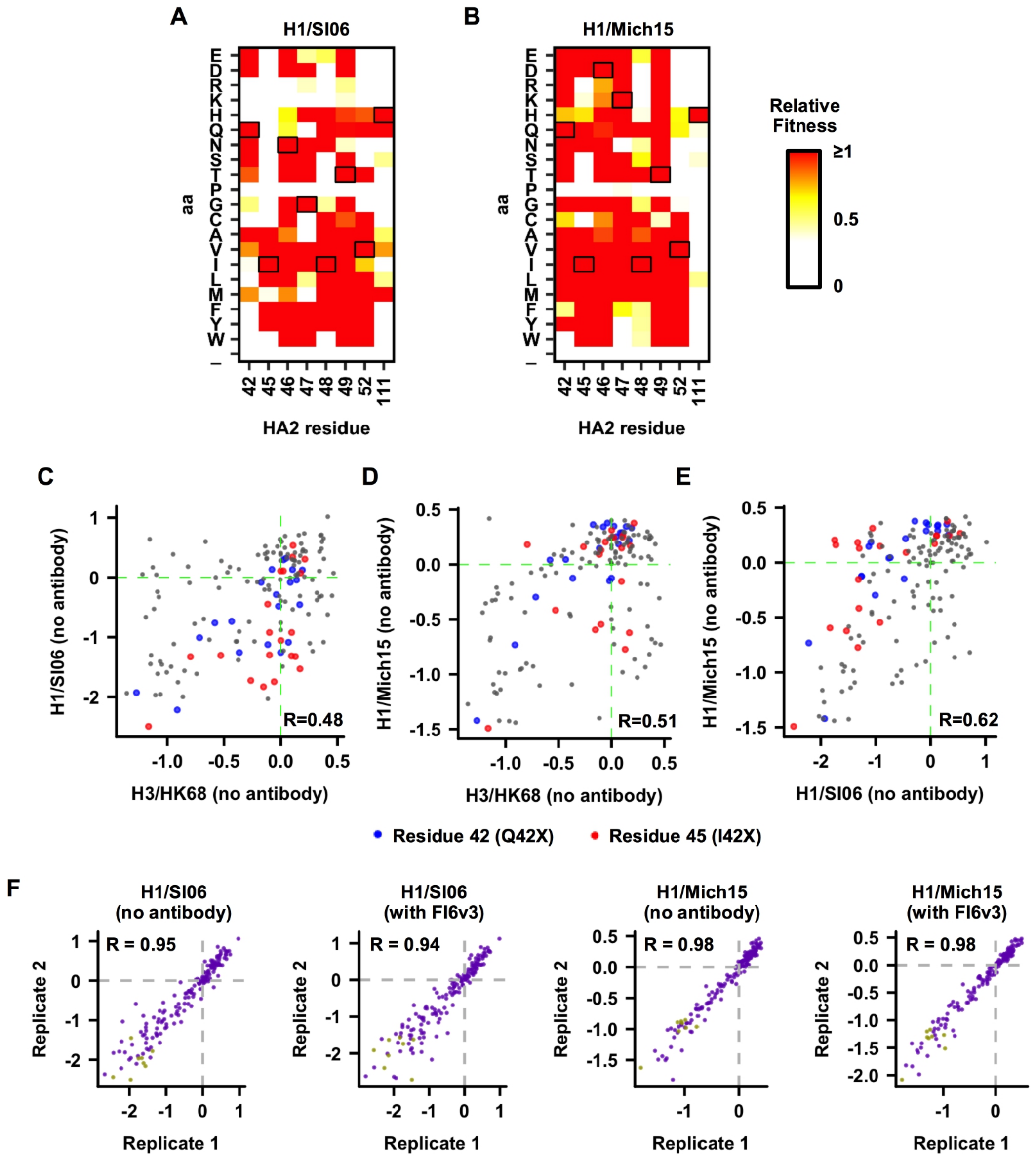


**Fig. S9. HA stem mutational fitness profiles of H1/SI06 and H1/Mich15. (A-B)** Based on the deep mutational scanning experiment, relative fitness of single mutants in **(A)** H1/SI06, and **(B)** H1/Mich15 viruses are shown. Relative fitness of wild type (WT) is set as 1. Residues correspond to WT sequence are boxed. **(C-E)** The $\log_{10}$ relative fitness of each single mutation in HA2 residues 42, 45, 46, 47, 48, 49, 52, and 111 of H3/HK68, H1/SI06, and H1/Mich15 in the absence of antibody are compared. **(C)** H3/HK68 vs H1/SI06. **(D)** H3/HK68 vs H1/Mich15. **(E)** H1/SI06 vs H1/Mich15. Each data point represents one mutant. Data points that represent mutations at residues 42 and 45 are colored in blue and red, respectively. Data points that represent mutations at other residues are colored in grey. Pearson correlation (R) is shown. These plots aim to analyze whether a give mutation has a similar fitness effect in different strains. For example, if a mutation that has a high fitness in one strain generally has a high fitness in another strain, a high correlation will be observed. **(F)** Relative fitness of each mutant in H1/SI06 and H1/Mich15 under different growth conditions is shown. Each data point represents one mutant. Pearson correlations (R) of the relative fitness of individual mutants between replicates are shown. Missense variants are colored in purple. Nonsense variants are colored in khaki green. The fitness measurements for each mutant in H1/SI06 and H1/Mich15 correlate well between biological replicates, demonstrating a high reproducibility of the results.

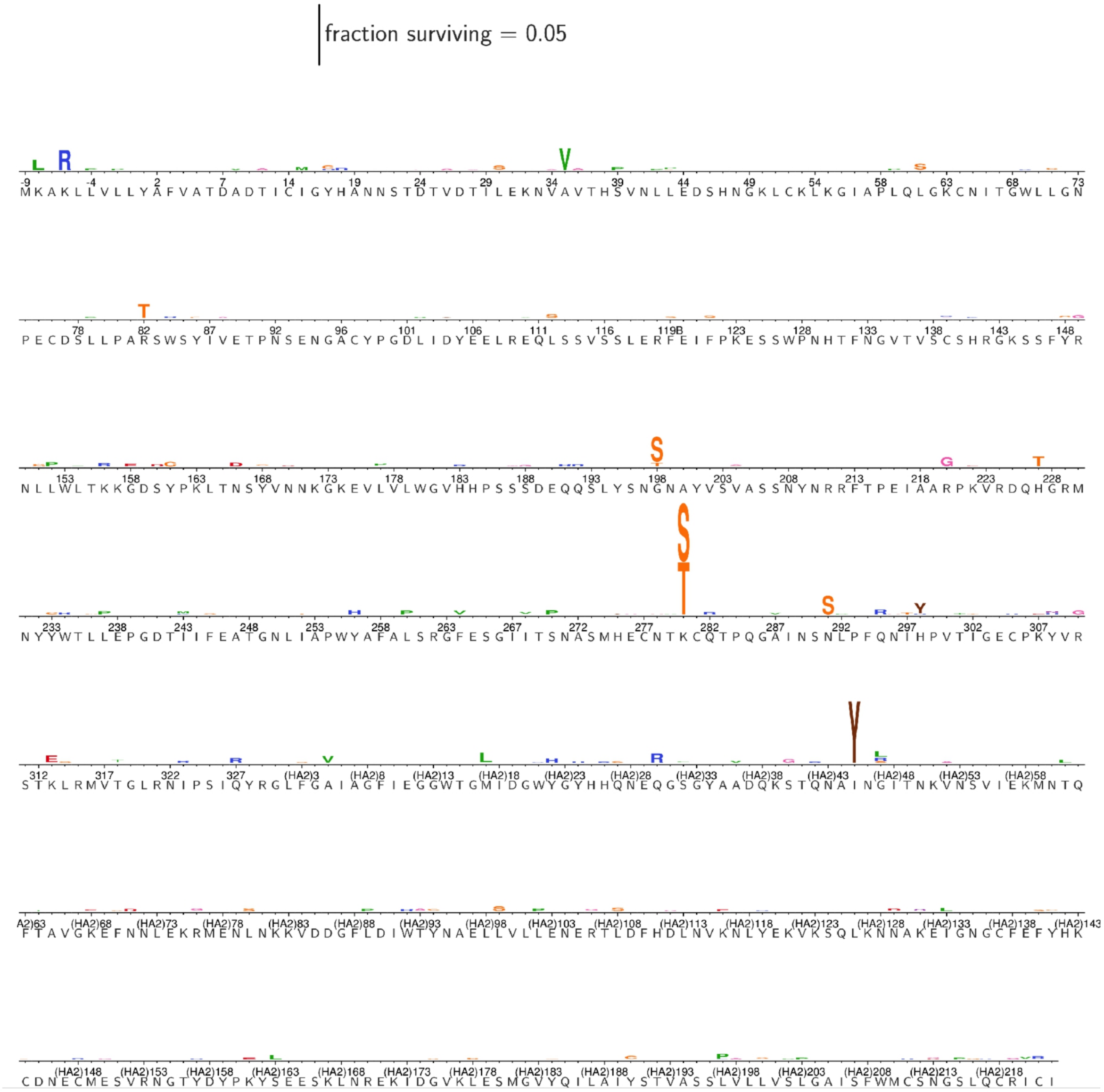


**Fig. S10. The excess fraction surviving selection with bnAb CR9114 for all amino-acid mutations in H1/WSN HA.** The height of each letter is proportional to the excess fraction of virion surviving with that mutation. The scale bar at the top of the plot relates the letter heights to the actual fractions. The sites are labeled using H3 numbering.

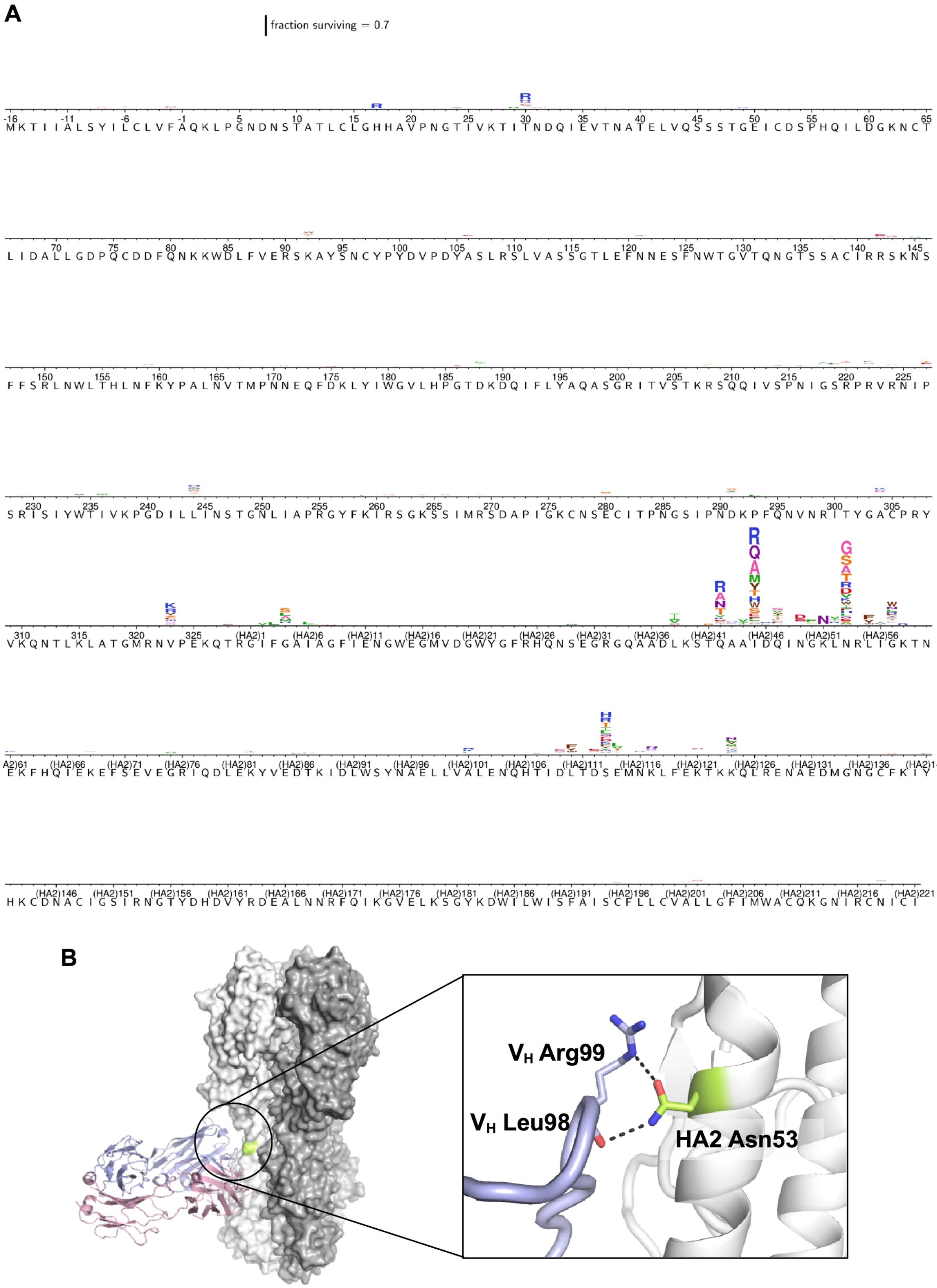


**Fig. S11. The excess fraction surviving selection with bnAb FI6v3 for all amino-acid**

**mutations in H3/Perth09 HA. (A)** The height of each letter is proportional to the excess fraction of virion surviving with that mutation. The scale bar at the top of the plot relates the letter heights to the actual fractions. The sites are labeled using H3 numbering. **(B)** HA2 Asn53 forms a hydrogen bond with the side chain of FI6v3 $V_H$ Arg99 as well as with the main-chain carbonyl of FI6v3 $V_H$ Leu98 (PDB 3ZTJ) (1). FI6v3 heavy chain is shown in blue, FI6v3 light chain in pink, and HA2 Asn53 in lime. Hydrogen bonds are represented by black dashed lines.

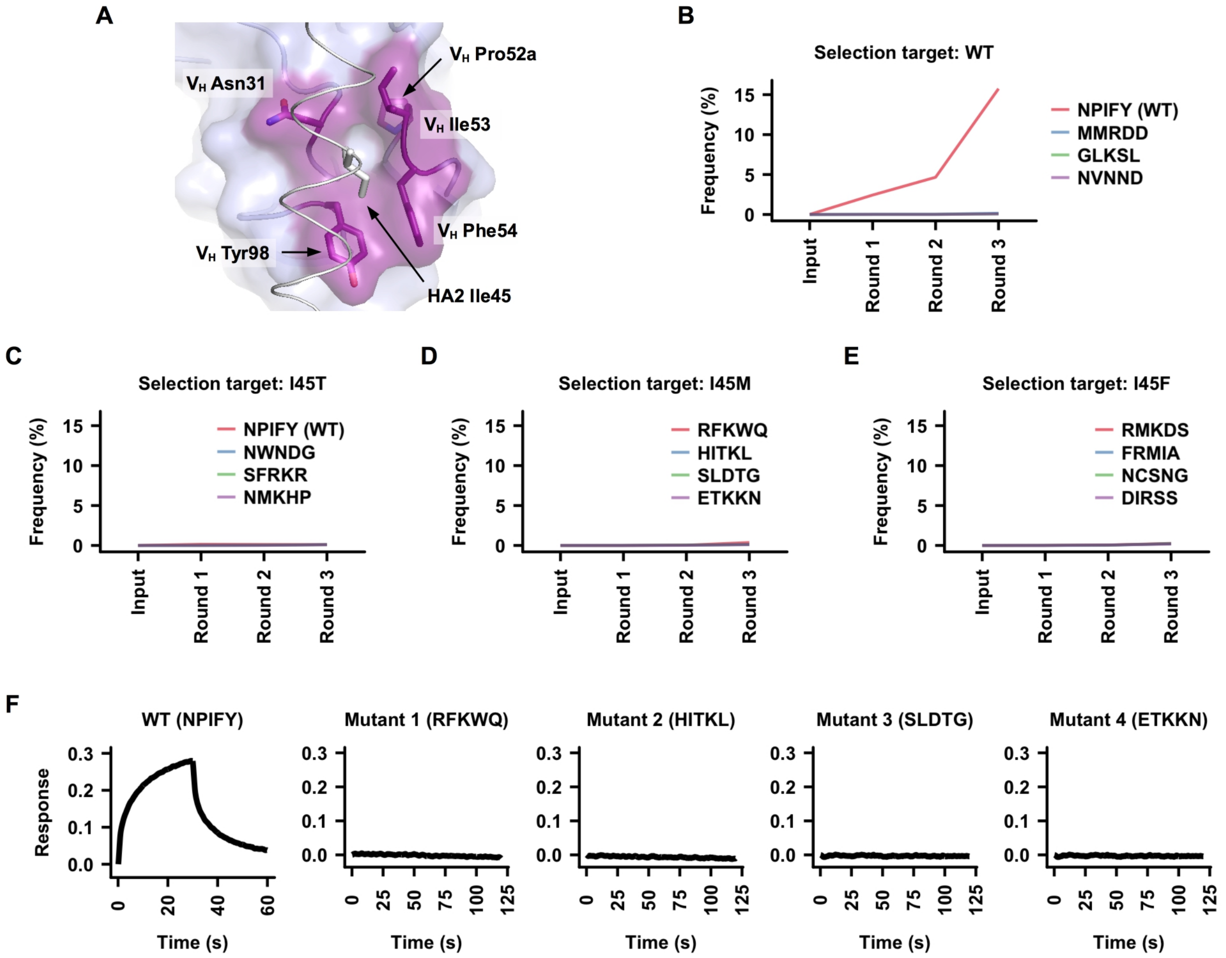


**Fig. S12. An attempt to overcome resistance mutations by evolving CR9114.** We tried to evolve CR9114 to bind to CR9114-resistance HA2 mutants using saturation mutagenesis and yeast display selection. **(A)** When CR9114 binds to HA, HA2-Ile45 is surrounded by multiple $V_H$ residues, including Asn31, Pro52a, Ile53, Phe54, and Tyr98. Saturation mutagenesis was performed at these five $V_H$ residues to generate a total of 3.2 million CR9114 variants ($20^5$ = 3,200,000). These CR9114 variants were displayed on the yeast surface in a Fab format (*56*) and underwent three rounds of selections against WT or mutant H3/HK68 HA. **(B-E)** The change in frequencies during the selection process were shown for: **(B)** the top four variants in post three-round selection against WT, **(C)** the top four CR9114 variants in post three-round selection against HA2-I45T mutant, **(D)** the top four CR9114 variants in post three-round selection against HA2-I45M mutant, and **(E)** the top four CR9114 variants in post three-round selection against HA2-I45F mutant. During the selection against WT H3/HK68 HA, the variant represents the WT CR9114 (NPIFY, abbreviated after the amino acids at the five residues of interest) was readily enriched. In contrast, none of the CR9114 variants showed significant enrichment during selections against those three HA2 mutants. This observation suggests that we failed to identify any variant from the mutant library that could overcome those three resistance HA2 mutants (I45T, I45M, and I45F). **(F)** The top four CR9114 variants in post three-round selection against HA2-I45M mutant, along with WT CR9114, were individually expressed and tested binding against recombinant HA2-I45M mutant. At 1 μM concentration, none of the CR9114 variants exhibited any binding to the recombinant HA2-I45M mutant, whereas WT CR9114 showed weak binding at the same concentration. This result demonstrates that it might be difficult to overcome those resistance mutants by simply introducing amino-acid substitutions in the bnAb.

**Table S1. X-ray data collection and refinement statistics**

| **Data collection** | H3/HK68 HA2-I45M | H3/HK68 HA2-I45T | H3/HK68 HA2-I45F |
|---|---|---|---|
| Beamline | SSRL 12-2 | SSRL 12-2 | SSRL 12-2 |
| Wavelength (Å) | 0.9795 | 0.9795 | 0.9795 |
| Space group | C2 | C2 | C2 |
| Unit cell parameters (Å and °) | a=208.9, b=131.0, c=72.6, β=98.4 | a=209.2, b=131.1, c=72.2, β=98.1 | a=208.4, b=131.4, c=72.3, β=98.1 |
| Resolution (Å) | 50-2.50 (2.60-2.50)[a] | 50-2.25 (2.35-2.25)[a] | 50-2.10 (2.20-2.10)[a] |
| Unique reflections | 65,790 (7,333)[a] | 89,621 (10,991)[a] | 110,282 (13,871)[a] |
| Redundancy | 5.9 (5.9)[a] | 5.0 (4.7)[a] | 5.6 (5.2)[a] |
| Completeness (%) | 97.6 (98.4)[a] | 97.4 (95.7)[a] | 97.4 (98.1)[a] |
| $\langle I/\sigma_I \rangle$ | 15.4 (1.5)[a] | 16.2 (1.3)[a] | 20.0 (1.7)[a] |
| $R_{sym}$[b] | 0.12 (0.87)[a] | 0.09 (0.94)[a] | 0.10 (0.83)[a] |
| $R_{pim}$[b] | 0.05 (0.39)[a] | 0.04 (0.49)[a] | 0.05 (0.40)[a] |
| $CC_{1/2}$[c] | 1.00 (0.70)[a] | 1.00 (0.68)[a] | 1.00 (0.77)[a] |
| $Z_a$[d] | 3 | 3 | 3 |
| **Refinement statistics** | | | |
| Resolution (Å) | 50-2.50 | 50-2.25 | 50-2.10 |
| Reflections (work) | 61,835 | 84,488 | 103,850 |
| Reflections (test) | 3,292 | 4,390 | 5,420 |
| $R_{cryst}$(%)[e] / $R_{free}$(%)[f] | 19.0 / 21.9 | 18.2 / 20.8 | 18.4 / 21.6 |
| No. of atoms | | | |
| Protein | 11,526 | 11,517 | 11,546 |
| Water | 321 | 666 | 846 |
| Glycan | 318 | 332 | 332 |
| Average *B*-value (Å$^2$) | | | |
| Protein | 57 | 50 | 48 |
| Water | 44 | 46 | 48 |
| Glycan | 88 | 79 | 71 |
| Wilson *B*-value (Å$^2$) | 48 | 38 | 32 |
| **RMSD from ideal geometry** | | | |
| Bond length (Å) | 0.011 | 0.011 | 0.011 |
| Bond angle (°) | 1.52 | 1.55 | 1.50 |
| **Ramachandran statistics (%)**[g] | | | |
| Favored | 96.5 | 97.1 | 97.1 |
| Outliers | 0.2 | 0.2 | 0.2 |
| **PDB code** | 6NHQ | 6NHP | 6NHR |

[a] Numbers in parentheses refer to the highest resolution shell.

[b] $R_{sym} = \Sigma_{hkl} \Sigma_i | I_{hkl,i} - \langle I_{hkl} \rangle | / \Sigma_{hkl} \Sigma_i I_{hkl,i}$ and $R_{pim} = \Sigma_{hkl} (1/(n-1))^{1/2} \Sigma_i | I_{hkl,i} - \langle I_{hkl} \rangle | / \Sigma_{hkl} \Sigma_i I_{hkl,i}$, where $I_{hkl,i}$ is the scaled intensity of the i$^{th}$ measurement of reflection h, k, l, $\langle I_{hkl} \rangle$ is the average intensity for that reflection, and n is the redundancy.

[c] $CC_{1/2}$ = Pearson correlation coefficient between two random half datasets.

[d] $Z_a$ is the number of HA protomers per crystallographic asymmetric unit.

[e] $R_{cryst} = \Sigma_{hkl} | F_o - F_c | / \Sigma_{hkl} | F_o | \times 100$, where $F_o$ and $F_c$ are the observed and calculated structure factors,

[f] $R_{free}$ was calculated as for $R_{cryst}$, but on a test set comprising 5% of the data excluded from refinement.

[g] Calculated with MolProbity (*61*).

**Table S2. Primers for construction of H3/HK68 mutant libraries**

| | |
|---|---|
| StemLib-VF | 5'-CGT ACG TCT CAA GTG CTT TTA AGA TCT GCT GCT TGT-3' |
| StemLib-VR | 5'-CGT ACG TCT CAT CGG AAA TGA ACA AGC TGT TTG AGA-3' |
| StemLib-WT-R | 5'-CGT ACG TCT CAC CGA GTC AGT CAG GTC AAT TGT ATG TTG ATT CT-3' |
| StemLib-WT-F | 5'-ACG TCT CAC ACT CAA GCA GCC ATC GAC CAA ATC AAT GGG AAA TTG AAC AGG GTA ATC GAG-3' |
| StemLib-111-R | 5'-CGT ACG TCT CAC CGA ATC SNN CAG GTC AAT TGT ATG TTG ATT CT-3' |
| StemLib-42-F | 5'-ACG TCT CAC ACT NNK GCC GCA ATC GAC CAA ATC AAT GGA AAA TTG AAC AGG GTA ATC GAG-3' |
| StemLib-45-F | 5'-ACG TCT CAC ACT CAA GCC GCA NNK GAC CAA ATC AAT GGC AAA TTG AAC AGG GTA ATC GAG-3' |
| StemLib-46-F | 5'-ACG TCT CAC ACT CAA GCC GCA ATC NNK CAA ATC AAT GGT AAA TTG AAC AGG GTA ATC GAG-3' |
| StemLib-47-F | 5'-ACG TCT CAC ACT CAA GCC GCT ATC GAC NNK ATC AAT GGA AAA TTG AAC AGG GTA ATC GAG-3' |
| StemLib-48-F | 5'-ACG TCT CAC ACT CAA GCC GCT ATC GAC CAA NNK AAT GGC AAA TTG AAC AGG GTA ATC GAG-3' |
| StemLib-49-F | 5'-ACG TCT CAC ACT CAA GCC GCT ATC GAC CAA ATC NNS GGT AAA TTG AAC AGG GTA ATC GAG-3' |
| StemLib-52-F | 5'-ACG TCT CAC ACT CAA GCC GCG ATC GAC CAA ATC AAT GGA AAA NNK AAC AGG GTA ATC GAG-3' |
| StemLib-42/45-F | 5'-ACG TCT CAC ACT NNK GCC GCG NNK GAC CAA ATC AAT GGC AAA TTG AAC AGG GTA ATC GAG-3' |
| StemLib-42/46-F | 5'-ACG TCT CAC ACT NNK GCC GCG ATC NNK CAA ATC AAT GGT AAA TTG AAC AGG GTA ATC GAG-3' |
| StemLib-42/47-F | 5'-ACG TCT CAC ACT NNK GCT GCA ATC GAC NNK ATC AAT GGA AAA TTG AAC AGG GTA ATC GAG-3' |
| StemLib-42/48-F | 5'-ACG TCT CAC ACT NNK GCT GCA ATC GAC CAA NNK AAT GGC AAA TTG AAC AGG GTA ATC GAG-3' |
| StemLib-42/49-F | 5'-ACG TCT CAC ACT NNK GCT GCA ATC GAC CAA ATC NNS GGT AAA TTG AAC AGG GTA ATC GAG-3' |
| StemLib-42/52-F | 5'-ACG TCT CAC ACT NNK GCT GCT ATC GAC CAA ATC AAT GGA AAA NNK AAC AGG GTA ATC GAG-3' |
| StemLib-45/46-F | 5'-ACG TCT CAC ACT CAA GCT GCT NNK NNK CAA ATC AAT GGC AAA TTG AAC AGG GTA ATC GAG-3' |
| StemLib-45/47-F | 5'-ACG TCT CAC ACT CAA GCT GCT NNK GAC NNK ATC AAT GGT AAA TTG AAC AGG GTA ATC GAG-3' |
| StemLib-45/48-F | 5'-ACG TCT CAC ACT CAA GCT GCG NNK GAC CAA NNK AAT GGA AAA TTG AAC AGG GTA ATC GAG-3' |
| StemLib-45/49-F | 5'-ACG TCT CAC ACT CAA GCT GCG NNK GAC CAA ATC NNS GGC AAA TTG AAC AGG GTA ATC GAG-3' |
| StemLib-45/52-F | 5'-ACG TCT CAC ACT CAA GCT GCG NNK GAC CAA ATC AAT GGT AAA NNK AAC AGG GTA ATC GAG-3' |
| StemLib-46/47-F | 5'-ACG TCT CAC ACT CAA GCG GCA ATC NNK NNK ATC AAT GGA AAA TTG AAC AGG GTA ATC GAG-3' |
| StemLib-46/48-F | 5'-ACG TCT CAC ACT CAA GCG GCA ATC NNK CAA NNK AAT GGC AAA TTG AAC AGG GTA ATC GAG-3' |
| StemLib-46/49-F | 5'-ACG TCT CAC ACT CAA GCG GCA ATC NNK CAA ATC NNS GGT AAA TTG AAC AGG GTA ATC GAG-3' |
| StemLib-46/52-F | 5'-ACG TCT CAC ACT CAA GCG GCT ATC NNK CAA ATC AAT GGA AAA NNK AAC AGG GTA ATC GAG-3' |
| StemLib-47/48-F | 5'-ACG TCT CAC ACT CAA GCG GCT ATC GAC NNK NNK AAT GGC AAA TTG AAC AGG GTA ATC GAG-3' |
| StemLib-47/49-F | 5'-ACG TCT CAC ACT CAA GCG GCT ATC GAC NNK ATC NNS GGT AAA TTG AAC AGG GTA ATC GAG-3' |
| StemLib-47/52-F | 5'-ACG TCT CAC ACT CAA GCG GCG ATC GAC NNK ATC AAT GGA AAA NNK AAC AGG GTA ATC GAG-3' |
| StemLib-48/49-F | 5'-ACG TCT CAC ACT CAA GCG GCG ATC GAC CAA NNK NNS GGC AAA TTG AAC AGG GTA ATC GAG-3' |
| StemLib-48/52-F | 5'-ACG TCT CAC ACT CAA GCG GCG ATC GAC CAA NNK AAT GGT AAA NNK AAC AGG GTA ATC GAG-3' |
| StemLib-49/52-F | 5'-ACG TCT CAC ACT CAA GCA GCA ATC GAC CAA ATC NNS GGA AAA NNK AAC AGG GTA ATC GAG-3' |

**Table S3. Primers for construction of H1/SI06 and H1/Mich15 mutant libraries**

| | | |
|---|---|---|
| H1/SI06 | StemLib-VF | 5'-CGT ACG TCT CAT CAA ATG TGA AGA ATC TGT ATG AGA-3' |
| | StemLib-VR | 5'-CGT ACG TCT CAT GTG CTT TTT TGG TCC GCA GCA TAG-3' |
| | StemLib-WT-F | 5'-ACG TCT CAC ACA CAA AAT GCC ATT AAC GGG ATT ACA AAC AAG GTG AAT TCT GTA ATC GAG-3' |
| | StemLib-42-F | 5'-ACG TCT CAC ACA NNK AAT GCC ATT AAC GGG ATT ACA AAC AAG GTG AAT TCT GTA ATC GAG-3' |
| | StemLib-45-F | 5'-ACG TCT CAC ACA CAA AAT GCC NNS AAC GGG ATT ACA AAC AAG GTG AAT TCT GTA ATC GAG-3' |
| | StemLib-46-F | 5'-ACG TCT CAC ACA CAA AAT GCC ATT NNK GGG ATT ACA AAC AAG GTG AAT TCT GTA ATC GAG-3' |
| | StemLib-47-F | 5'-ACG TCT CAC ACA CAA AAT GCC ATT AAC NNK ATT ACA AAC AAG GTG AAT TCT GTA ATC GAG-3' |
| | StemLib-48-F | 5'-ACG TCT CAC ACA CAA AAT GCC ATT AAC GGG NNS ACA AAC AAG GTG AAT TCT GTA ATC GAG-3' |
| | StemLib-49-F | 5'-ACG TCT CAC ACA CAA AAT GCC ATT AAC GGG ATT NNK AAC AAG GTG AAT TCT GTA ATC GAG-3' |
| | StemLib-52-F | 5'-ACG TCT CAC ACA CAA AAT GCC ATT AAC GGG ATT ACA AAC AAG NNK AAT TCT GTA ATC GAG-3' |
| | StemLib-WT-R | 5'-CGT ACG TCT CAT TGA GTC ATG AAA ATC CAA AGT CCT CTC ATT TT-3' |
| | StemLib-111-R | 5'-CGT ACG TCT CAT TGA GTC SNN AAA ATC CAA AGT CCT CTC ATT TT-3' |
| H1/Mich15 | StemLib-VF | 5'-CGT ACG TCT CAT CAA ATG TGA AGA ACT TGT ATG AAA-3' |
| | StemLib-VR | 5'-CGT ACG TCT CAT GTG CTC TTC AGG TCG GCT GCA TAT-3' |
| | StemLib-WT-F | 5'-ACG TCT CAC ACA CAA AAT GCC ATT GAC AAG ATT ACT AAC AAA GTA AAT TCT GTT ATT GAA-3' |
| | StemLib-42-F | 5'-ACG TCT CAC ACA NNK AAT GCC ATT GAC AAG ATT ACT AAC AAA GTA AAT TCT GTT ATT GAA-3' |
| | StemLib-45-F | 5'-ACG TCT CAC ACA CAA AAT GCC NNS GAC AAG ATT ACT AAC AAA GTA AAT TCT GTT ATT GAA-3' |
| | StemLib-46-F | 5'-ACG TCT CAC ACA CAA AAT GCC ATT NNK AAG ATT ACT AAC AAA GTA AAT TCT GTT ATT GAA-3' |
| | StemLib-47-F | 5'-ACG TCT CAC ACA CAA AAT GCC ATT GAC NNK ATT ACT AAC AAA GTA AAT TCT GTT ATT GAA-3' |
| | StemLib-48-F | 5'-ACG TCT CAC ACA CAA AAT GCC ATT GAC AAG NNS ACT AAC AAA GTA AAT TCT GTT ATT GAA-3' |
| | StemLib-49-F | 5'-ACG TCT CAC ACA CAA AAT GCC ATT GAC AAG ATT NNS AAC AAA GTA AAT TCT GTT ATT GAA-3' |
| | StemLib-52-F | 5'-ACG TCT CAC ACA CAA AAT GCC ATT GAC AAG ATT ACT AAC AAA NNK AAT TCT GTT ATT GAA-3' |
| | StemLib-WT-R | 5'-CGT ACG TCT CAT TGA ATC GTG ATA GTC CAA AGT TCT TTC ATT TT-3' |
| | StemLib-111-R | 5'-CGT ACG TCT CAT TGA ATC SNN ATA GTC CAA AGT TCT TTC ATT TT-3' |
